# Rational Design and Synthesis of Selective PRMT4 Inhibitors: a New Chemotype for Development of Cancer Therapeutics

**DOI:** 10.1101/2020.11.17.387233

**Authors:** Mathew Sutherland, Alice Li, Anissa Kaghad, Dimitrios Panagopoulos, Fengling Li, Magdalena Szewczyk, David Smil, Cora Scholten, Léa Bouché, Timo Stellfeld, Cheryl H. Arrowsmith, Dalia Barsyte, Masoud Vedadi, Ingo V. Hartung, Holger Steuber, Robert Britton, Vijayaratnam Santhakumar

## Abstract

Protein arginine N-methyl transferase 4 (PRMT4) asymmetrically dimethylates arginine residues of histone H3 and non-histone proteins. The overexpression of PRMT4 in several cancers has stimulated interest in the discovery of inhibitors as biological tools and potentially therapeutics. While several PRMT4 inhibitors have been reported, most display poor selectivity against other members of the PRMT family of methyl transferases. Here, we report the structure-based design of a new class of alanine containing 3-arylindoles as potent and selective PRMT4 inhibitors and describe key structure activity relationships for this class of compounds.

---

Arginine methyl transferases catalyze both symmetric and asymmetric methylation of arginine residues in the histone H3 proteins using the methyl group from S-adenosyl-L-methionine (SAM).^1^ These methyl transferases regulate a variety of biological processes including transcriptional activation,^2^ RNA splicing,^3^ cell cycle regulation,^4^ DNA damage response,^5^ and cell differentiation,^6^ while also catalyzing the methylation of a variety of non-histone proteins.^7^ Type I arginine methyl transferases (PRMTs), PRMT1, 2, 3, 4, 6, and 8, catalyze mono and asymmetric dimethylation. The Type II PRMTs PRMT5 and 9 catalyze mono and symmetric di-methylation while PRMT7 catalyzes mono methylation of arginines.^8^

PRMT4 has been implicated in several malignancies and is highly overexpressed in ~75% of colorectal cancers^9^ as well as in prostate carcinoma and androgen-independent prostate carcinoma.^10^ PRMT4 is a critical factor in the pathway of estrogen-stimulated breast cancer growth^11^ and its overexpression is associated with poor prognosis in this disease.^12^ Knock-out studies in breast cancer cell lines show that PRMT4 regulates breast cancer cell migration and metastasis.^13^ Moreover, pharmacological inhibition of PRMT4 with selective inhibitors is effective in reducing the growth of multiple myeloma cell lines^14^ as well as in vivo mouse models of multiple myeloma^15^.

The majority of reported PRMT4 inhibitors shows moderate to poor selectivity against other Type I PRMTs^16,17^ and/or lack of cellular activity,^18–20^ with the notable exceptions of PRMT4 selective chemical probes 1 and 2 reported by Structural Genomics Consortium^14,21^ and Epizyme,^15^ respectively. Here, we report the development of indole based, potent and PRMT4 selective inhibitors starting from a dual PRMT4/6 inhibitor scaffold. Notably, we relied on the co-crystal structure of a hit compound 3a (Figure 1) with PRMT6 and molecular modeling with reported PRMT4 structures to design PRMT4 selective inhibitors and identify key structural features relevant to PRMT4 selectivity.

**Figure 1.**
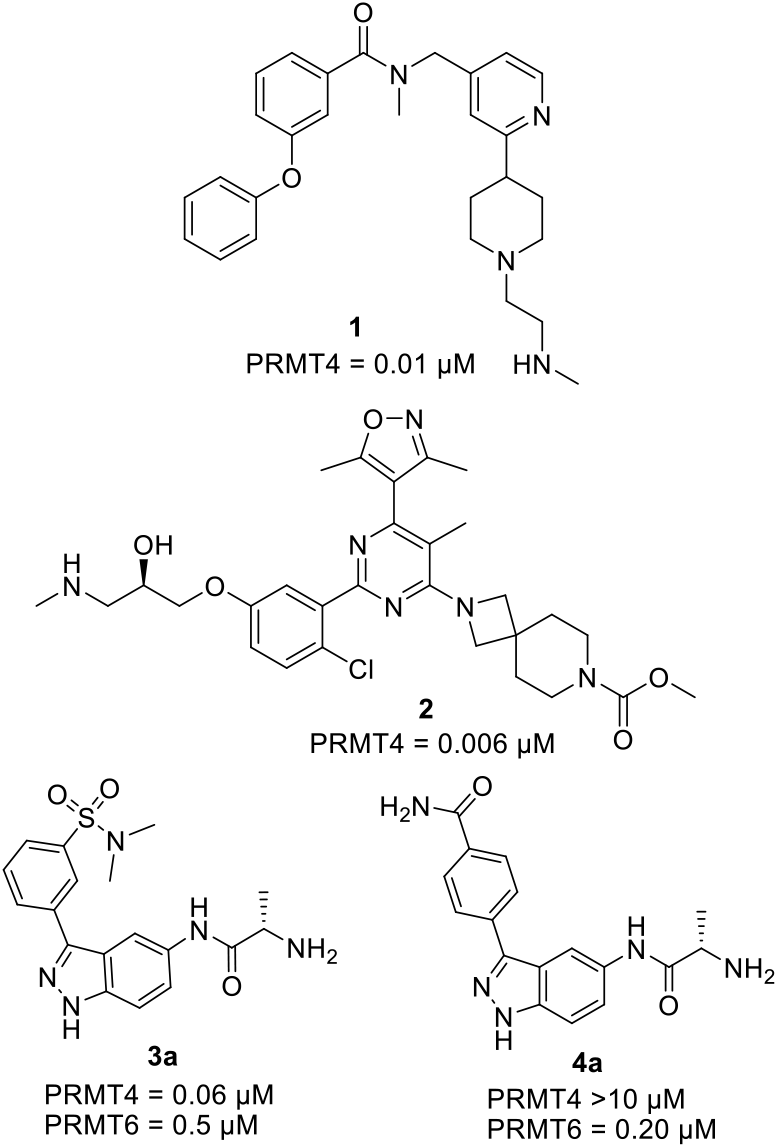
PRMT4 inhibitors **1** and **2** reported by the Structural Genomics Consortium and Epizyme, respectively, and dual PRMT4/6 inhibitors identified through screening campaigns

Owing to its potent PRMT4 inhibitory activity (IC_50_ 0.06 μM) and moderate selectivity over PRMT6 family (8-fold), the indazole **3a** (entry 1, Table 1) was selected as a starting point for the development of a selective PRMT4 inhibitor. Unfortunately, attempts to co-crystallize **3a** or the structurally related indazole **4a** (Figure 1) with PRMT4 were unsuccessful. However, we were able to obtain the co-crystal structure of **3a** with PRMT6 (Figure 2A). Based on this structural insight, superimposition of the PRMT6-bound structure of indazole **3a** in the reported PRMT4 structure^19^ (Figure 2B) suggested that the corresponding indole may improve selectivity for PRMT4. Specifically, in the PRMT6-bound structure of **3a**, H-bonding between Glu59 and the indazole NH was identified as a key binding interaction. Thus, it was proposed that decreasing the acidity of the N-H from indazole (pKa ~ 14) to indole (pKa ~21) would attenuate interactions with PRMT6. Additionally, the reduced polar surface area (PSA) of the corresponding indole would expectedly result in improved hydrophobic interactions with PRMT4 (Figure 2B). To test this hypothesis, indole **4^b^** was synthesized following the synthetic sequence described in Scheme 1. PRMT4 and PRMT6 activity assays were preformed according to the published procedures^14^. and we were pleased to find that this compound showed a moderate loss in PRMT6 activity and coincident gain in PRMT4 activity compared to **4a** (Table 1, entry 2 and 3). From a small collection of indazoles, it was noted that replacement of the amide with a *meta*-sulfonamide (entries 1, 2) resulted in improved PRMT4 activity. Inspired by these observations, the corresponding *meta*-methyl sulfone analogue of indole **4b** was prepared (entry 4), resulting in further improved PRMT4 activity and selectivity. Notably, the *meta*-methyl sulfone **3b** (entry 4) proved to be 197-fold selective against PRMT6 (PRMT4 IC_50_ = 40 nM) and >50-fold selective against PRMT1,3,5-9. Docking studies of the sulfonamide **3a** in both PRMT4 and PRMT6 suggested that the improved PRMT4 activity is likely due to interactions within the unique hydrophobic binding pocket in PRMT4 (Figure 2B). This additional subpocket in PRMT4 is mainly created by Gln 149 and Phe 153, while the corresponding less space-demanding residues Leu 46 and Cys 50 do not generate s similar pocket in PRMT6.

**Figure 2.**
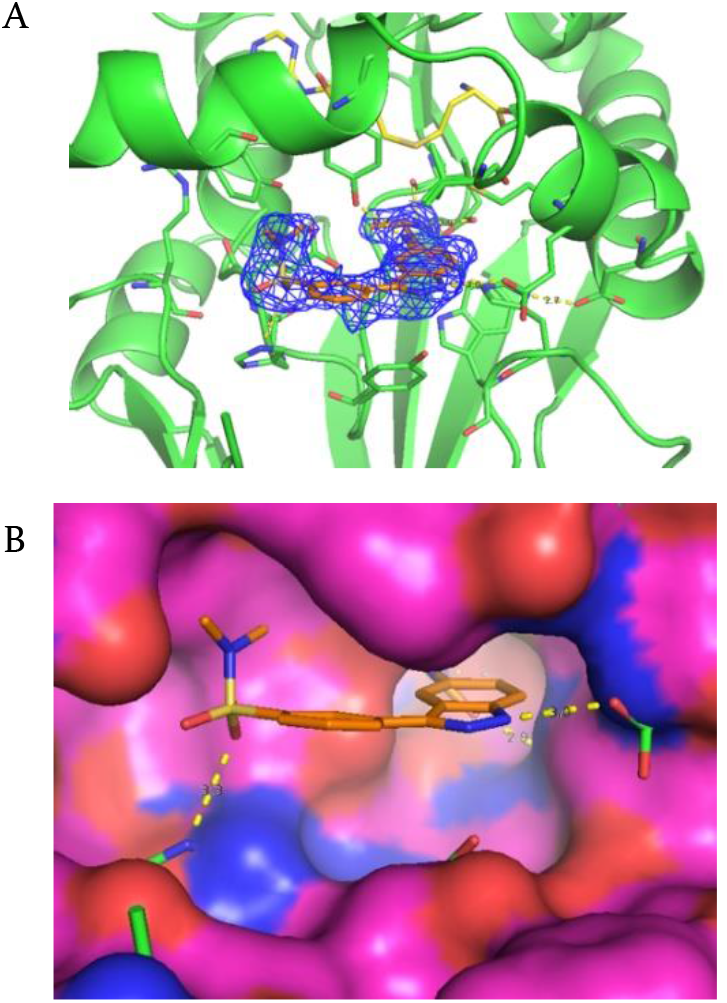
(A) The PRMT6-bound structure of **3a**; (B) PRMT6-bound conformation of **3a** superimposed in binding site of PRMT4 highlights a hydrophobic pocket (occupied by sulfonamide group) not present in PRMT6.

**Table 1:**
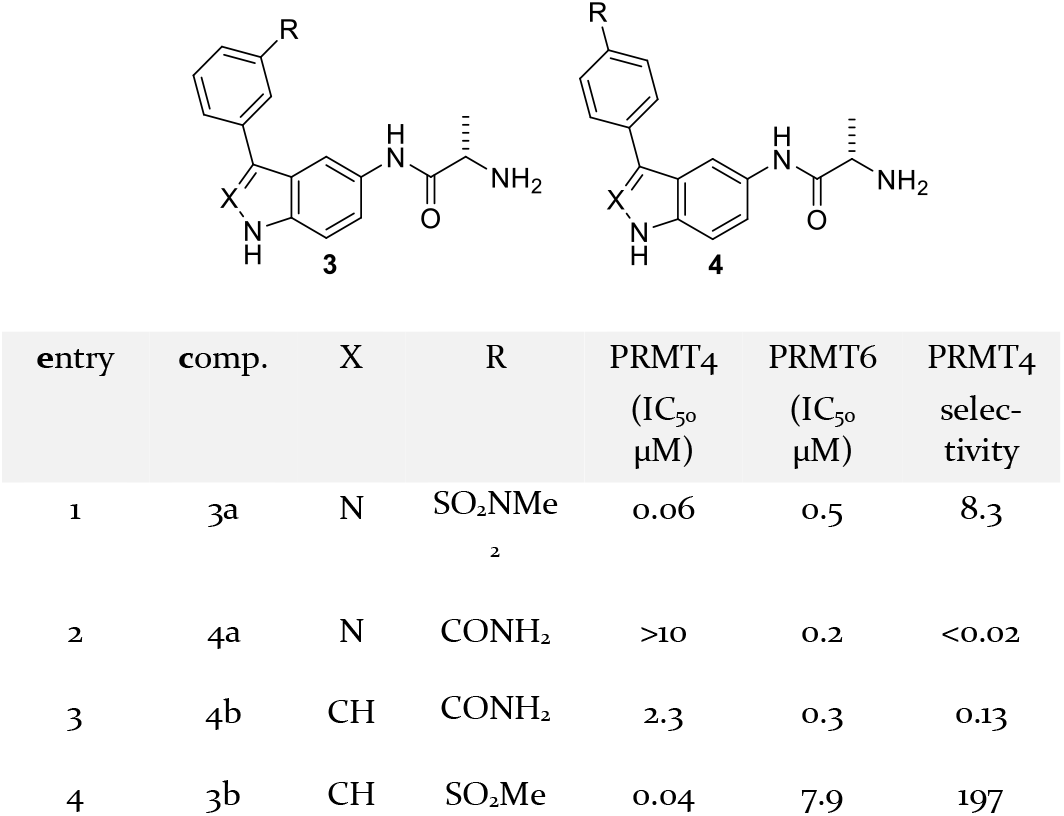
PRMT4 and PRMT6 activity of indazoles and indoles 3 and 4.

**Scheme 1.**
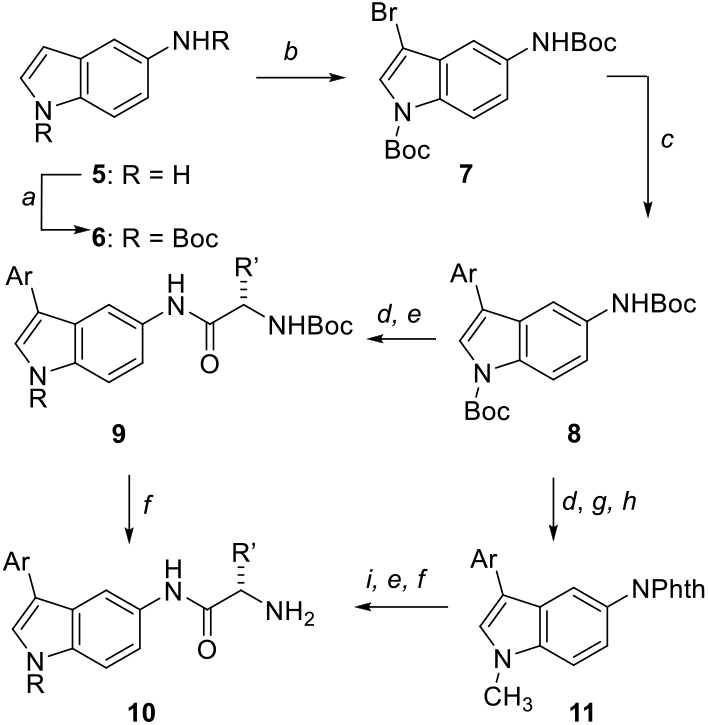
General synthetic route for preparing indole-based PRMT4 inhibitors. Reagents and conditions: (a) Boc_2_O, THF; (b) NBS, THF; (c) ArB(OR)_2_, K_2_CO_3_, Pd(PPh_3_)_4_ or Pd(dppf)Cl_2_•CH_2_Cl_2_, THF:H_2_O (3:1), 80 °C; (d) TFA; (e) N-Boc-amino acid, DIPEA, pyBOP, DMF (f) TFA; g) phthalic anhydride, toluene, reflux; h) K_2_CO_3_, CH3I, DMF, rt; i) H_2_NNH_2_, MeOH,

To further probe the effect of modifications at the meta-position in indole 3b, a series of sulfones and sulfonamides was prepared in a straightforward manner as summarized in Scheme 1 and Table 2. Here, we found that exchanging the methyl sulfone for a dimethyl sulfonamide (e.g. compound 12, entries 1 and 2) resulted in a 2.5-fold gain in PRMT4 activity (IC_50_ = 0.014 μM) and a 3-fold improvement in selectivity (650-fold). This result indicated that a more lipophilic dimethyl sulfonamide better exploits the hydrophobic binding pocket in the PRMT4 active site. However, this modification was accompanied by a loss in potency when more sterically hindered sulfonamides were examined. For example, the diethyl sulfonamide 13 (entry 3, IC_50_ = 3.5 μM) and cyclic sulfonamide 14 (entry 4, IC_50_ = 0.89 μM) proved to be less active against PRMT4. While the smaller five membered ring sulfonamide 15 (entry 5) was tolerated (IC_50_ = 0.18 μM), this compound was still less potent and selective than original methyl sulfone 3b. Several analogues of the methyl sulfone 3b were also synthesized and it was found that the isopropyl sulfone 16 (entry 6) was a potent PRMT4 inhibitor (IC_50_ = 0.03 μM). Here again, a similar trend to that seen with sulfonamides was observed. Specifically, increasing the size of the alkyl sulfone led to a significant loss in potency (e.g., 17; IC_50_ = 3.3 μM). This data suggested that the hydrophobic binding pocket in PRMT4 could not accommodate groups larger than the dimethyl sulfonamide or isopropyl sulfone. As a result, dimethyl sulfonamide 12 (entry 2) and isopropyl sulfone 16 (entry 6) were selected as the lead molecules for further optimization.

**Table 2.** Biological evaluation of sulfone and sulfonamide series

| Entry | Comp. | R | PRMT4<br>(IC<sub>50</sub> μM) |
| --- | --- | --- | --- |
| 1 | 3b | Me | 0.04 |
| 2 | 12 | NMe<sub>2</sub> | 0.01 |
| 3 | 13 | NEt<sub>2</sub> | 3.5 |
| 4 | 14 | N(CH<sub>2</sub>)<sub>5</sub> | 0.89 |
| 5 | 15 | N(CH<sub>2</sub>)<sub>4</sub> | 0.18 |
| 6 | 16 | iPr<sub>2</sub> | 0.01 |
| 7 | 17 | C(CH<sub>2</sub>)<sub>4</sub> | 3.3 |

Having identified that both the sulfone and sulfonamide confer excellent PRMT4 activity and selectivity, we then turned our attention towards the amino acid side of the molecule. Here, we aimed to increase lipophilicity to improve cellular permeability and perhaps potency and selectivity. With this in mind, we chose the methyl sulfone as our starting point for the amino acid structure activity relationship study. We probed the size of the amino acid with the L-proline analogue **18a** (entry 1, Table 3) and found this compound was not active (PRMT4 IC_50_ >10 μM). The ethyl amine and *N*-methyl ethyl amine **19a** and **19b**, respectively, were also synthesized based on the common use of ethyl amine in PRMT inhibitors^16,17,22^. Unfortunately, in both cases we observed a significant loss in potency (PRMT4 IC_50_ >10 μM in both cases). A similar amino acid SAR study on the isopropyl sulfone and dimethyl sulfon-amide scaffolds was undertaken through the synthesis of compounds **18b-18c** and **18d-18h**, respectively (entries 4-11, Table 3). Here, we examined methylation and incorporation of an azetidine for the isopropyl sulfone and in the case of the sulfonamide, we investigated incorporation of an azetidine, methylation, glycine incorporation, “symmetric” dimethylation and cyclopropanation. In the case of the isopropyl sulfone, each modification resulted in a decrease in PRMT4 activity (PRMT4 IC_50_ = 0.6 to >10 μM). In general, the dimethyl sulfonamide analogues 18d-18h were more potent. In particular, the glycine analogue 18f proved to be a low nM inhibitor of PRMT4 (IC_50_ = 0.01 μM).

**Table 3.** SAR of the amino acid moiety

| Entry | Comp. | R<sub>1</sub> | R<sub>2</sub> | R<sub>3</sub> | PRMT<sub>4</sub><br>(IC<sub>50</sub> μM) |
| --- | --- | --- | --- | --- | --- |
| 1 | 18a | Me | (CH<sub>2</sub>)<sub>3</sub> | (CH<sub>2</sub>)<sub>3</sub> | >10 |
| 2 | 19a | Me | N/A | H | >10 |
| 3 | 19b | Me | N/A | Me | 27 |
| 4 | 18b | iPr<sub>2</sub> | Me | Me | >10 |
| 5 | 18c | iPr<sub>2</sub> | (CH<sub>2</sub>)<sub>2</sub> | (CH<sub>2</sub>)<sub>2</sub> | 0.6 |
| 7 | 18d | NMe<sub>2</sub> | (CH<sub>2</sub>)<sub>2</sub> | (CH<sub>2</sub>)<sub>2</sub> | 0.11 |
| 8 | 18e | NMe<sub>2</sub> | Me | Me | 4.2 |
| 9 | 18f | NMe<sub>2</sub> | H | H | 0.01 |
| 10 | 18g | NMe<sub>2</sub> | Me<sub>2</sub> | H | 0.33 |
| 11 | 18h | NMe<sub>2</sub> | (CH<sub>2</sub>)<sub>2</sub> | H | >10 |

At this point, we examined the cell permeability of the most promising compounds 3b, 12, 16 and 18d. The results of the Caco-2 assay are summarized in Figure 3. From the series methyl sulfone 3b, dimethyl sulfonamide 12 and isopropyl sulfone 16, the dimethyl sulfonamide proved to be the most permeable (2.13 nm/s for 12 vs 0.23 nm/s for 3b). Disappointingly, replacement of alanine for azetidine in an effort to reduce H-bond donors, increase lipophilicity and enhance cellular permeability of the compounds increased the efflux ratio by ~3 fold (Figure 3). Based on this data we further explored a series of analogues that incorporated the dimethyl sulfonamide core.

**Figure 3.**
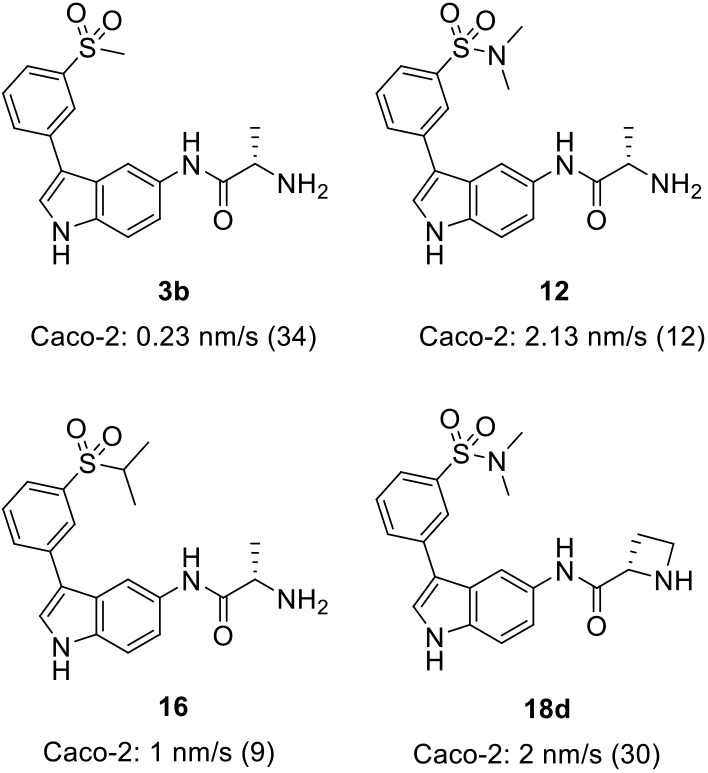
Caco-2 results for key compounds (efflux ratio in parenthesis)

As the co-crystal structure of indazole 3a bound to PRMT6 indicated that the aryl ring was largely solvent exposed, we next focused on modifications aimed to increase lipophilicity using the most potent dimethylsulfonamide core. Thus, analogues 20-22 were synthesized as summarized in Scheme 1. Of this series, the N-Me indole 20 proved to be the most potent and selective compound (PRMT4 IC_50_ = <2 nM, >500-fold selectivity against PRMT6, Table 4, entry 1). In order to assess the effect of attenuating the basicity of the free amine on membrane permeability, the fluoroalanine 21 was synthesized^2^3 (entry 2, Table 4), which unfortunately proved to be ~10-fold less potent and selective. To further reduce the total polar surface area of the molecule, we examined the dimethyl amide 22, however, this analogue proved also to be less active and selective. In summary, the N-methylated derivative 20 is the most potent compound of our series against PRMT4, indicating that H-bonding of the indole-NH to Asn 162 is by far less relevant for PRMT4 binding, while it is important for the charge-reinforced interaction to Glu 59 in PRMT6. Hence, methylation of the indole-N is a key driver to achieve selectivity against PRMT6.

**Table 4.** Evaluation of methylated indole derivatives

| Entry | Comp. | R | R' | PRMT<sub>4</sub><br>(IC<sub>50</sub> μM) |
| --- | --- | --- | --- | --- |
| 1 | 20 | SO<sub>2</sub>NMe<sub>2</sub> | Me | <0.002 |
| 2 | 21 | SO<sub>2</sub>NMe<sub>2</sub> | CH<sub>2</sub>F | 0.18 |
| 3 | 22 | CONMe<sub>2</sub> | Me | 0.22 |

## Chemistry

The general preparation of compounds described herein was performed as shown in Scheme 1, starting from commercially available 5-aminoindole **5**. Boc Protection of the amine and indole nitrogen atoms^24^ gave the bis-Boc-protected compound **6**, which was exposed to NBS to furnish the corresponding 3-brominated product **7**^25^. This later material then engaged in a Suzuki reaction^26^ with a suitable *meta*-substituted sulfone or sulfonamide derivative. Following the Suzuki reaction, global deprotection was accomplished by treatment with trifluoroacetic acid. The 5-amino group was then coupled to a Boc-protected amino acid using pyBOP coupling conditions^27,28^. Finally, deprotection of the amino acid using trifluoroacetic acid yielded the desired indole analogues as their corresponding trifluoroacetate salts. In the case of analogues **20-22**, prior to amino acid coupling, the 5-amino group was protected as a phthalyl group and the indole nitrogen was methylated using methyl iodide and potassium carbonate (**11**, Scheme 1). Deprotection of the phthalimide was effected by treatment with hydrazine and the free amine was then carried through a similar sequence of steps as described above (i.e., peptide coupling and Boc-deprotection).

## Evaluation of cellular activity of PRMT4 inhibitors

PRMT4 has been shown to methylate BAF155 at R1064^13^. We have evaluated key compounds **3b**, **20** and **22**, however, none of the compounds showed significant reduction in BAF155 methylation when tested up to 100 μM (48 h exposure in HEK293 cells), while methylation of BAF155 was abrogated by 2 μM of the PRMT4 selective inhibitor TP-64 (Figure 4). The absence of any significant cellular activity of compound **3b** can be attributed to its poor permeability. Even though compounds **20** and **22** were expected to show enhanced permeability because of reduced H-bond donors (**20** and **22**) and reduced polar surface area (**22**), the absence of cellular activity indicates these changes were not sufficient to increase the cellular permeability of these series of compounds.

**Figure 4.**
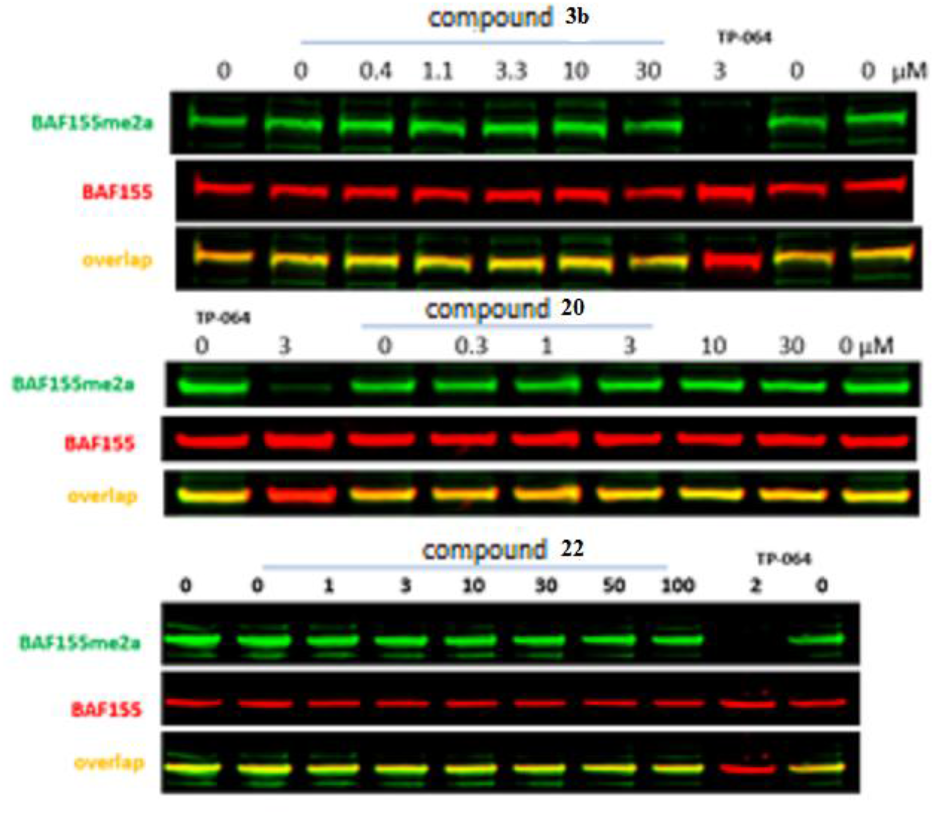
Effect of PRMT4 dependent BAF155 asymmetric dimethylation in HEK293T cells upon treatment with compounds 3b, 20 and 22 up to 100 μM over 2 days.

In conclusion, we report the design, synthesis and evaluation of a new series of 3-arylindole alanine-based PRMT4 inhibitors that are both potent and selective over the closely related PRMT6. We demonstrate that selectivity for PRMT4 can be achieved by carefully tuning the substitution pattern of the aromatic ring and that amino acid substitutions are not well-tolerated. Furthermore, methylation of the indole nitrogen resulted in a potent and selective in-vitro PRMT4 inhibitor. Despite these efforts, none of the compounds described herein achieved on-target effects in cells. Nonetheless, these indoles represent a new chemotype for further development of cell active PRMT4 inhibitors, which will deepen our understanding of the intricate biology of PRMT4 and may engender the design of new anti-tumor PRMT4-selective inhibitors. These compounds could possibly be used as handles to develop PROTACs to degrade PRMT4 selectively. Linker attachment and E3 antagonist components used for developing PROTACs will likely influence the cellular permeability of the PROTAC compounds and therefore the poor cellular permeability of our PRMT4 inhibitors are not likely to be a hinderance for cell active PROTAC development.

## General Chemistry

All reagents and starting materials were purchased from Sigma Aldrich, TCI, Alfa Aesar, CarboSynth, and AK Sci and were used without further purification. Dichloromethane was distilled from CaH_2_ and stored under nitrogen, THF was distilled from sodium wire/benzophenone ketyl radical and stored under nitrogen. Column chromatography was carried out with 230-400 mesh silica gel (E. Merck, Silica Gel 60). Concentration and removal of trace solvents was done via a Buchi rotary evaporator using acetone-dry-ice condenser and a Welch vacuum pump. Nuclear magnetic resonance (NMR) spectra were recorded using deuterochloroform (CDCl_3_), deuteromethanol (CD_3_OD), deuteroacetonitrile (CD_3_CN) or deuterodimethyl sulfoxide (DMSO-d_6_) as the solvent. Signal positions (δ) are given in parts per million from tetramethylsilane (δ 0) and were measured relative to the signal of the solvent (^1^H NMR: CDCl_3_: δ 7.26; CD_3_OD: δ3.31; CD_3_CN: δ1.96; DMSO-dβ: δ 2.50; ^13^C NMR: CDCl_3_: δ 77.16; CD_3_OD: δ 49.00; CD_3_CN: δ1.32; DMSO-d_6_: 39.5). Coupling constants (*J* values) are given in Hertz (Hz) and are reported to the nearest 0.1 Hz. ^1^H NMR spectral data are tabulated in the order: multiplicity (s, singlet; d, doublet; t, triplet; q, quartet; sept, septet; m, multiplet; br, broad), coupling constants, number of protons. NMR spectra were recorded on a Bruker Avance 600 equipped with a QNP or TCI cryoprobe (600 MHz), Bruker 400 (400 MHz) or Bruker 500 (500 MHz). High performance liquid chromatography (HPLC) analysis was performed on an Agilent 1100 HPLC, equipped with a variable wavelength UV-Vis detector. High-resolution mass spectrometry was performed on an Agilent 6210 TOF LC/MS.

## General Procedures

### General Procedure A: Suzuki-Miyaura Coupling

A pressure vial charged with a stir bar, bromoindole **7** (1.0 equiv.), boronic acid or ester (1.0 – 1.6 equiv.), K_2_CO_3_ (3.0 equiv.), and Pd(PPh_3_)_4_ or Pd(dppf)Cl_2_.CH_2_Cl_2_ (0.10 equiv.) was placed under vacuum and then filled with nitrogen. A mixture of degassed THF and H_2_O (0.09 M THF-H_2_O 3:1 unless otherwise indicated) was then added and the resulting mixture was stirred under an atmosphere of nitrogen at 80 °C for 18 hours or until the reaction was complete as monitored by TLC analysis. The reaction mixture was then cooled to room temperature and concentrated under reduced pressure. The residue was then dissolved in EtOAc and washed with saturated aqueous NaHCO_3_ and brine. The organic layer was dried over MgSO_4_ and concentrated to afford the crude aryl-indole product. Purification of the crude product by flash chromatography (silica gel, Et_2_O or EtOAc and hexanes) afforded the pure coupled product.

### General Procedure B: Amide Coupling and Deprotection

To a stirred solution of the aryl-indole intermediate (1.0 equiv.) in dry dimethylformamide (DMF) (0.1 M) at room temperature was added *N,N*-diisopropylethylamine (DIPEA) (5 equiv.), followed by benzotriazol-1-yl-oxytripyrrolidinophosphonium hexafluorophosphate (PyBOP) (1-2 equiv.) and the protected amino acid (1-2 equiv.). The resulting solution was stirred at room temperature until completion of the reaction as monitored by TLC. The reaction mixture was then diluted with saturated aqueous NaHCO_3_ and extracted with EtOAc (3 times). The combined organic layers were washed with saturated aqueous LiCl (3 times), dried over MgSO_4_, and concentrated to afford the crude product (brown gum), which was used directly in the next step without further purification. The crude coupled product was dissolved in TFA (neat, 0.1 M) and stirred at room temperature until the reaction was complete as monitored by TLC analysis. Purification of the crude indole by RP-HPLC (using a SiliCycle SiliaChrom dtC18 semi-preparative column (5 μm, 100Å, 10 x 250 mm) with a flow rate of 5 mL/min eluting with solvent (A: 0.1 % TFA in H_2_O B: 0.1 % TFA in ACN) on gradients of 2 → 30%, or 2 → 100% solvent B over 15 minutes as indicated afforded the final compounds.

### General Procedure C: Synthesis of Sulfonamides

To a stirred solution of amine (1.05 equiv.) in dry pyridine (0.2 M) at 0 °C was added dropwise (for liquids) or in small portions (for solids) the sulfonyl chloride (1.0 equiv.). The reaction mixture was warmed to room temperature and stirred until the reaction was complete as monitored by TLC analysis. The reaction mixture was then concentrated under reduced pressure and the residue was dissolved in EtOAc and washed with 0.5 M HCl (2 times). The organic layer was then dried over MgSO_4_, filtered and concentrated to afford the sulfonamide. The sulfonamide was used in subsequent reactions without further purification.

### General Procedure D: Boronic Ester Synthesis

A flask was charged with a stir bar, aryl bromide (1.0 equiv.), B_2_pin_2_ (1.0 equiv.), NaHCO_3_ (2.50 equiv.), and Pd(ddpf)Cl_2_ (0.05 equiv.). The flask was then placed under vacuum and filled with nitrogen. Degassed DMSO (0.2 M) was added to the reaction vessel, and the reaction mixture was stirred under an atmosphere of nitrogen at 80 °C for 18 hours or until the reaction was complete as monitored by TLC analysis. The reaction mixture was then cooled to room temperature and diluted with equal parts H_2_O and EtOAc, then filtered through Celite and the Celite was rinsed with EtOAc. The filtrate was then washed with H_2_O and brine, dried over MgSO_4_ and concentrated to afford the crude product. Purification of the crude product by flash chromatography (silica gel, Et_2_O or EtOAc and hexanes) afforded the aryl-boronic ester.

### General Procedure E: Sulfone Synthesis from Thiophenol Precursors

A stirred solution of substituted thiophenol (1 equiv.), K_2_CO_3_ (1.4 equiv.), and secondary bromoalkane (1.2 equiv.) in dry acetone (0.3 M), was stirred under nitrogen at reflux until completion of the reaction was observed by TLC (*ca*. 18 hours). The reaction mixture was cooled to room temperature, diluted with H_2_O, and extracted with Et_2_O (3 times). The combined organic layers were washed with brine, dried over MgSO_4_, and concentrated to afford the crude aryl thioether intermediate. To a stirred solution of the crude aryl thioether intermediate (1.0 equiv.) in MeOH (0.16 M) at 0 °C was added oxone (potassium peroxymonosulfate) (3.0 equiv.) in H_2_O (0.5 M). The resulting white suspension was warmed to room temperature over 2 hours and stirred at room temperature until completion was observed by TLC. The reaction mixture was then diluted with H_2_O and extracted with EtOAc (2 times). The combined organic layers were washed with brine, dried over MgSO_4_, and concentrated to afford the aryl sulfone. The aryl sulfone was used in subsequent reactions without further purification.

### General Procedure F: Phthalimide protection

To a stirred solution of aminoindole intermediate (1.0 equiv.) in toluene (0.2 M) was added phthalic anhydride (1.3 equiv.). The resultant solution was heated to reflux until completion of the reaction was observed by TLC. The reaction mixture was then cooled down to room temperature and concentrated under reduced pressure to afford a crude product which was used without further purification unless otherwise indicated.

### General Procedure G: *N*-methylation of indole scaffolds

To a stirred solution of protected indole intermediate (1.0 equiv.) in dry dimethylformamide (DMF) (0.2 M) was added potassium carbonate (5 equiv.) followed by methyl iodide (3 equiv.). The resultant solution was stirred at 60 °C until completion of the reaction was observed by TLC. The reaction mixture was then diluted with water and extracted with EtOAc (3 times). The combined organic layers were washed with aqueous saturated LiCl (3 times), dried over MgSO_4_, and concentrated to afford the crude methylated product, which was used directly in the next step without further purification unless otherwise indicated.

### General procedure H: Phthalimide deprotection

To a stirred solution of *N*-methylindole intermediate (1.0 equiv.) in methanol (0.06 M) was added hydrazine hydrate (1.3 equiv.). The resultant solution was stirred at room temperature until completion of the reaction was observed by TLC. The reaction mixture was then concentrated, diluted in dichloromethane and filtered. The resulting filtrate was concentrated under reduced pressure to afford the amino indole product, which was used directly in the next step without further purification.

### General procedure I: *Boc*-deprotection

A solution of Boc-protected intermediate (1 eq) in TFA (0.1 M) was stirred at room temperature until completion of the reaction was observed by TLC. Concentration of the reaction mixture under reduced pressure afforded the deprotected aryl-indole product which was used without further purification unless otherwise indicated.

### tert-butyl (S)-(1-((3-bromo-1H-indazol-5-yl)amino)-1-oxopropan-2-yl)carbamate (23)

To a solution of (*tert*-butoxycarbonyl)-L-alanine (1.78 g, 9.43 mmol) in DMF (15 ml) at 21 °C was added HATU (4.30 g, 11.32 mmol) and *N,N*-diisopropylethylamine (4.93 ml, 28.29 mmol). The resulting solution was stirred for 10 min prior to the addition of 3-bromo-1*H*-indazol-5-amine (2.00 g, 9.43 mmol), and subsequent stirring for a further 16 h. The mixture was diluted with brine (100 mL), the aqueous layer extracted with EtOAc (3 x 50 mL), and the combined organic extracts dried with anhydrous sodium sulfate before being filtered and concentrated under reduced pressure. The residue was purified by column chromatography on silica gel (Biotage SNAP 25 g column, 0-20% EtOAc/hexanes as the eluent, 25 CV) to afford **23** as a light yellow solid (2.53 g, 70%). **^1^H NMR** (400 MHz, DMSO) δ (ppm) = 13.34 (s, 1H), 10.03 (s, 1H), 8.03 (s, 1H), 7.49-7.55 (m, 2 H), 7.07 (br. d, *J* = 6.8 Hz, 1 H), 4.13 (br. t, *J* ≈ 6.8 Hz, 1 H), 2.28 (br d, *J* = 7.3 Hz, 3 H), 1.39 (s, 9 H); LRMS = 386.2 [M+H]^+^.

### (S)-2-amino-N-(3-(3-(N,N-dimethylsulfamoyl)phenyl)-1H-indazol-5-yl)propenamide (3a)

To a solution of **23** (57.5 mg, 0.15 mmol) in 1-methyl-2-pyrrolidinone (NMP) (1 mL), were added subsequently [3-(dimethylsulfamoyl)phenyl]boronic acid (68.7 mg, 0.30 mmol, dissolved in 0.53 mL 1-methyl-2-pyrrolidinone),[1,1’-Bis(diphenylphosphino)ferrocene]dichloropalladium(II), complex with dichloromethane (24.5 mg, 0.03 mmol, dissolved in 1 mL NMP) and potassium carbonate (62.2 mg, 0.45 mmol, dissolved in water 0.5 mL). The reaction mixture was heated to 100 °C and shaken for 24 h. The crude mixture was filtered through a pad of activated MP Alumina N (EcoChrom TM) and washed with NMP and concentrated under vacuum. The residue was dissolved in a TFA/CH_2_Cl_2_ mixture (1:1, 2 mL) and shaken for 24 h and finally dried using a Christ-centrifuge to give 4.62 mg of the title compound (7% yield). LC-MS method: Instrument MS: Waters ZQ; Instrument HPLC: Waters UPLC Acquity; column: Acquity BEH C_18_ (Waters), 50 mm x 2.1mm, 1.7 μm; Solvent: A: 0.1% formic acid in water, solvent B: MeCN; Gradient: 0.0min 99% A – 1.6 min 1% A – 1.8 min 1%A – 1.81 min 99% A – 2.0 min 99% A; oven: 60 °C; flow: 0.800 mL/min; UV-Detection PDA 210-400nm. R_t_ = 0.83 min; LRMS (ESI): m/z = 388 [M+H]^+^

### (S)-4-(5-(2-aminopropanamido)-1H-indazol-3-yl)benzamideSynthesis of benzamide (4a)

Carbamate **23** (57.5 mg, 0.15 mmol, dissolved in 1 mL NMP), (4-carbamoylphenyl)boronic acid (49.5 mg, 0.30 mmol, dissolved in 0.53 mL NMP) and [1,1’-Bis(diphenylphosphino)ferrocene]dichloropalladium(II), complex with dichloromethane (24.5 mg, 0.03 mmol, dissolved in 1 mL NMP) along with potassium carbonate (62.2 mg, 0.45 mmol, dissolved in water 0.5 mL) were heated to 100 °C and shaken for 24 h. The microtitre plates (MTPs) were dried by Zirbus-centrifuge and then were dissolved again in 2 mL of a TFA/acetonitrile (1:1) mixture. The reaction mixture was further shaken for 1d at room temperature. The MTPs were dried again and 2 mL NMP were added. The precipitated material was filtered off and purified by preparative HPLC to give 4.01 mg of the title compound (8% yield). LC-MS method: Instrument MS: Waters ZQ; Instrument HPLC: Waters UPLC Acquity; column: Acquity BEH C_18_ (Waters), 50 mm x 2.1mm, 1.7 μm; Solvent A: 0.1% formic acid in water, solvent B: MeCN; Gradient: 0.0min 99% A – 1.6 min 1% A – 1.8 min 1%A – 1.81 min 99% A – 2.0 min 99% A; oven: 60 °C; flow: 0.800 mL/min; UV-Detection PDA 210-400 nm. R_t_ = 0.50 min; LRMS (ESI): m/z = 324 [M+H]^+^

### tert-butyl 5-((tert-butoxycarbonyl)amino)-1H-indole-1-carboxylate (6)

To a stirred brown slurry of 5-aminoindole (1.00 g, 7.56 mmol, 1.0 equiv.) in THF (38 mL, 0.2 M) was added triethylamine (1.05 mL, 7.56 mmol, 1.0 equiv.), Boc_2_O (3.30 g, 15.1 mmol, 2.0 equiv.), and DMAP (1.39 g, 11.3 mmol, 1.5 equiv.). The resulting slurry was stirred at room temperature under a nitrogen atmosphere (not sealed) for 2 days. The reaction mixture was diluted with *ca*. 60 mL of 1.0 M aqueous HCl and extracted with EtOAc (2 times). The combined organic layers were dried over MgSO_4_ and concentrated under reduced pressure to afford the crude *bis*-protected aminoindole species. Purification of the crude product (silica gel, Et_2_O:Hexanes 1:9) provided the *bis*-protected aminoindole 6 (1.30 g, 52 %). **^1^ NMR**: (400 MHz, CDCl_3_) δ (ppm) = 8.02 (d, *J* = 8.8 Hz, 1H), 7.74 (s, 1H), 7.56 (d, *J* = 3.7 Hz, 1H), 7.13 (d, *J* = 9.0 Hz, 1H), 6.65 – 6.54 (m, 1H), 6.49 (d, *J* = 3.6 Hz, 1H), 1.66 (s, 9H), 1.53 (s, 9H). **^13^C NMR**: (100 MHz, CDCl_3_) δ (ppm) = 153.3, 149.8, 133.7, 131.6, 131.2, 126.7, 116.4, 115.4, 110.9, 107.5, 83.7, 80.4, 28.5, 28.3. **HRMS**: (ESI) *m/z* calculated for C_18_H_24_N_2_O_4_ [2M+H]^+^ 655.3545, found 655.3574.

### tert-butyl 3-bromo-5-((tert-butoxycarbonyl)amino)-1H-indole-1-carboxylate (7)

To a stirred solution of **6** (1.52 g, 4.56 mmol, 1.0 equiv.) in THF (13.4 mL, 0.34 M) was added NBS (852 mg, 4.79 mmol, 1.05 equiv.). The reaction vessel was wrapped in foil to exclude light and stirred at room temperature for 18 hours, after which it was concentrated under reduced pressure. The reaction residue was then dissolved in Et2O and filtered to remove the white precipitate. The filtrate was washed with saturated aqueous sodium metabisulfite, saturated aqueous NaHCO_3_, water, brine, dried over MgSO_4_, filtered, and concentrated under reduced pressure to afford **7** (1.46 g, 77 %). **^1^H NMR**: (500 MHz, CDCl_3_) δ (ppm) = 8.03 (s, 1H), 7.64 (s, 1H), 7.61 (s, 1H), 7.25 (d, *J* = 8.0 Hz, 1H), 6.60 (s, 1H), 1.65 (s, 9H), 1.54 (s, 9H). **^13^C NMR**: (125 MHz, CDCl3) δ (ppm) = 153.1, 148.9, 134.5, 131.0, 130.1, 125.6, 117.5, 115.7, 109.2, 98.0, 84.4, 80.7, 28.5, 28.3. **HRMS**: (ESI) *m/z* calculated for C_18_H_23_BrN_2_O_4_ [M+NH_4_]^1^ 428.1179, found 428.1204.

### 3-(3-(methylsulfonyl)phenyl)-1H-indol-5-amine (24)

The title compound was prepared according to general procedure A using **7** (583 mg, 1.5 mmol), 4,4,5,5-tetramethyl-2-(3-(methylsulfonyl)phenyl)-1,3,2-dioxaborolane (400 mg, 1.5 mmol), K_2_CO_3_ (588 mg, 4.3 mmol), Pd(PPh_3_)_4_ (164 mg, 0.15 mmol), degassed THF (12 mL) and degassed water (4 mL). Purification by column chromatography afforded the protected coupled product, which was subsequently deprotected in TFA (15 mL) and concentrated under reduced pressure to afford the TFA salt of **24** as a brown solid (453 mg, 80%). **^1^H NMR**: (500 MHz, CD_3_OD) δ (ppm) = 8.21 (dd, *J* = 1.8, 1.7 Hz, 1H), 8.00 (ddd, *J* = 7.8, 1.8, 1.4 Hz, 1H), 7.91 (d, *J* = 2.1 Hz, 1H), 7.85 (ddd, *J* = 7.9, 1.7, 1.4 Hz, 1H), 7.81 (s, 1H), 7.71 (dd, *J* = 7.9, 7.8 Hz, 1H), 7.63 (d, *J* = 8.6 Hz, 1H), 7.22 (dd, *J* = 8.6, 2.1 Hz, 1H), 3.19 (s, 3H). **^13^C NMR**: (125 MHz, CD_3_OD) δ (ppm) = 142.8, 138.3, 138.2, 133.1, 131.2, 127.5, 126.8, 126.1, 125.5, 124.6, 117.5, 116.8, 114.6, 114.2, 44.4. **HRMS**: (ESI) *m/z* calculated for C15H14N2O_2_S [M+H]^+^ 287.0849, found 287.0857.

### (S)-2-amino-N-(3-(3-(methylsulfonyl)phenyl)-1H-indol-5-yl)propenamide (3b)

The title compound was prepared according to general procedure B using the aryl-indole **24** (63.2 mg, 0.158 mmol), (*tert*-butoxycarbonyl)*-L*-alanine (60 mg, 0.32 mmol), PyBOP (165 mg, 0.32 mmol), DIPEA (0.21 mL, 0.79 mmol), and dry DMF (1.6 mL). Purification by RP-HPLC (using a SiliCycle SiliaChrom dtC18 semipreparative column (5 μm, 100Å, 10 x 250 mm) with a flow rate of 5 mL/min eluting with solvent (A: 0.1 % TFA in water B: 0.1 % TFA in MeCN) on a gradient of (2 → 100) % solvent B over 15 min, tR = 5.68 min) of the crude deprotected product afforded the TFA salt of **3b** as a colorless solid (11 mg, 18%). **^1^H NMR**: (600 MHz, CD_3_OD) δ (ppm) = 8.24 (d, *J* = 1.9 Hz, 1H), 8.22 (s, 1H), 7.99 (d, *J* = 7.7 Hz, 1H), 7.80 (d, *J* = 7.6 Hz, 1H), 7.70 – 7.64 (m, 2H), 7.45 (d, *J* = 8.7 Hz, 1H), 7.33 (dd, *J* = 8.7, 2.0 Hz, 1H), 4.10 (q, *J* = 7.0 Hz, 1H), 3.20 (s, 3H), 1.63 (d, *J* = 7.1 Hz, 3H). **^13^C NMR**: (150 MHz, CD_3_OD) δ (ppm) = 169.1, 142.5, 139.1, 136.2, 132.9, 132.1, 131.0, 126.4, 126.0, 126.0, 124.9, 117.2, 116.4, 113.2, 111.8, 50.9, 44.5, 17.8. **HRMS**: (ESI) *m/z* calculated for C_18_H_19_N_3_O_3_S [M+H]^+^ 358.1220, found 358.1230.

### 4-(5-amino-1H-indol-3-yl)benzamide (25)

The title compound was prepared according to general procedure A using **7** (454 mg, 1.10 mmol), *para*-carbamoylphenyl-boronic acid (291 mg, 1.76 mmol), K_2_CO_3_ (457 mg, 3.31 mmol), Pd(ddpf)Cl_2_.CH_2_Cl_2_ (90 mg, 0.11 mmol), degassed THF/water (3:1, 20 mL, 0.056 M). Purification by column chromatography afforded the protected coupled product, which was subsequently deprotected in TFA (11 mL) and concentrated under reduced pressure to afford the TFA salt of **25** as a brown solid (231 mg, 55%). **^1^H NMR**: (500 MHz, CD_3_OD) δ (ppm) = 7.98 (d, *J* = 8.4 Hz, 2H), 7.92 (d, *J* = 2.1 Hz, 1H), 7.80 – 7.74 (m, 3H), 7.61 (d, *J* = 8.6 Hz, 1H), 7.19 (dd, *J* = 8.6, 2.1 Hz, 1H). ^13^**C NMR**: (125 MHz, CD_3_OD) δ (ppm) = 172.2, 140.5, 138.2, 132.0, 129.4, 127.7, 127.3, 127.0, 124.5, 117.6, 117.2, 114.43, 114.39. **HRMS**: (ESI) *m/z* calculated for C_15_H_13_N_3_O [M+H]^+^ 252.1131, found 252.1144.

### (S)-4-(5-(2-aminopropanamido)-1H-indol-3-yl)benzamide (4b)

The title compound was prepared according to general procedure B using the aryl-indole **25** (25 mg, 0.068 mmol), (*tert*-butoxycarbonyl)*-L*-alanine (17 mg, 0.072 mmol), PyBOP (43 mg, 0.082 mmol), DIPEA (0.06 mL, 0.34 mmol), and dry DMF (0.68 mL). Purification by RP-HPLC (using a SiliCycle SiliaChrom dtC18 semipreparative column (5 μm, 100Å, 10 x 250 mm) with a flow rate of 5 mL/min eluting with solvent (A: 0.1 % TFA in water B: 0.1 % TFA in MeCN) on a gradient of (2 → 100) % solvent B over 15 min, tR = 4.52 min) of the crude deprotected product afforded the TFA salt of **4b** as a colorless solid (9 mg, 32%). **^1^H NMR**: (500 MHz, CD_3_OD) δ (ppm) = 8.23 (d, *J* = 1.7 Hz, 1H), 7.94 (d, *J* = 8.5 Hz, 2H), 7.77 (d, *J* = 8.5 Hz, 2H), 7.64 (s, 1H), 7.43 (d, *J* = 8.7 Hz, 1H), 7.32 (dd, *J* = 8.7, 1.7 Hz, 1H), 4.09 (q, *J* = 7.1 Hz, 1H), 1.64 (d, *J* = 7.1 Hz, 3H). **^13^C NMR**: (125 MHz, CD_3_OD) δ (ppm) = 172.4, 169.0, 141.4, 136.3, 131.8, 131.4, 129.3, 127.5, 126.6, 126.0, 117.3, 117.2, 113.1, 112.5, 50.9, 17.7. **HRMS**: (ESI) *m/z* calculated for C_18_H_18_N_4_O_2_ [M+H]^+^ 323.1503, found 323.1477.

### 3-bromo-N,N-dimethylbenzenesulfonamide (26)

The title compound was prepared according to general procedure C, using dimethylamine (0.024 mL, 0.36 mmol), dry pyridine (1.8 mL), and 3-bromobenzenesulfonyl chloride (0.056 mL, 0.39 mmol). Workup of the reaction mixture as detailed in general procedure C afforded the aryl-sulfonamide **26** as a colorless oil (87 mg, 95 %). **^1^H NMR**: (500 MHz, CDCl_3_) δ (ppm) = 7.73 (dd, *J* = 1.9, 1.7 Hz, 1H), 7.71 (ddd, *J* = 8.0, 1.9, 1.0 Hz, 1H), 7.43 (ddd, *J* = 7.9, 1.7, 1.0 Hz, 1H), 7.43 (dd, *J* = 8.0, 7.9 Hz, 1H), 2.74 (s, 6H)

### N,N-dimethyl-3-(4,4,5,5-tetramethyl-1,3,2-dioxaborolan-2-yl)benzenesulfonamide (27)

The title compound was prepared according to general procedure D, using aryl-sulfonamide **26** (71 mg, 0.27 mmol), B2Pin2 (205 mg, 0.81 mmol), NaOAc (88 mg, 1.08 mmol) and Pd(dppf)Cl_2_ (22 mg, 0.027 mmol) in degassed dioxane (1.1 mL). Purification of the crude material by column chromatography (EtOAc:hexanes 10:90) afforded the 3-pinacolboronate aryl-sulfonamide **27** as a colorless solid (53 mg, 63 %). **^1^H NMR**: (500 MHz, CDCl_3_) δ (ppm) = 8.20 (dd, *J* = 1.9, 1.1 Hz, 1H), 8.01 (ddd, *J* = 7.5, 1.5, 1.1 Hz, 1H), 7.86 (ddd, *J* = 7.9, 1.9, 1.5 Hz, 1H), 7.37 (dd, *J* = 7.9, 7.5 Hz, 1H), 2.72 (s, 6H), 1.35 (s, 12H).

### tert-butyl 5-((tert-butoxycarbonyl)amino)-3-(3-(N,N-dimethylsulfamoyl)phenyl)-1H-indole-1-carboxylate (28)

The title compound was prepared according to general procedure A using **7** (49.5 mg, 0.12 mmol), 3-pinacolboronate aryl-sulfonamide **27** (37 mg, 0.12 mmol), K_2_CO_3_ (50 mg, 0.36 mmol), Pd(PPh_3_)_4_ (14 mg, 0.012 mmol), degassed THF (1.0 mL) and degassed water (0.35 mL). Purification by column chromatography (EtOAc:hexanes 10:90) afforded **28** as a pale yellow solid (32 mg, 52%). **^1^H NMR**: (500 MHz, CDCl_3_) δ (ppm) = 8.12 (d, *J* = 8.5 Hz, 1H), 8.03 (br s, 2H), 7.85 (dt, *J* = 7.8, 1.2 Hz, 1H), 7.76 (br d, *J* = 7.6 Hz, 2H), 7.64 (t, *J* = 7.8, 1H), 7.17 (dd, *J* = 8.9, 1.8 Hz, 1H), 6.56 (br s, 1H), 2.81 (s, 6H), 1.69 (s, 9H), 1.51 (s, 9H).

### (S)-2-amino-N-(3-(3-(N,N-dimethylsulfamoyl)phenyl)-1H-indol-5-yl)propenamide (12)

The title compound was prepared according to the general procedures I and B using the Boc-protected-aryl-indole **28** (55 mg, 0.106 mmol) and TFA (1.0 mL) followed by treatment with (*tert*-butoxycarbonyl)*-L*-alanine (24 mg, 0.13 mmol), PyBOP (66 mg, 0.53 mmol), DIPEA (0.13 mL, 0.13 mmol), and dry DMF (1.0 mL). Final Boc-deprotection of the crude mixture using TFA (1.0 mL) and subsequent purification by RP-HPLC (using a SiliCycle SiliaChrom dtC18 semipreparative column (5 μm, 100À, 10 x 250 mm) with a flow rate of 5 mL/min eluting with solvent (A: 0.1 % TFA in water B: 0.1 % TFA in MeCN) on a gradient of (2 → 100) %) solvent B over 15 min, t_R_ = 6.2 min) afforded the TFA salt of **12** as a colorless solid (8 mg, 19%). **^1^H NMR**: (400 MHz, CD_3_OD) δ (ppm) = 8.32 (d, *J* = 1.9 Hz, 1H), 8.06 (br s, 1H), 7.95 (dt, *J* = 6.7, 2.1 Hz, 1H), 7.68-7.61 (m, 3H), 7.45 (d, *J* = 8.8 Hz, 1H), 7.26 (dd, *J* = 8.8, 2.1 Hz, 1H), 4.08 (q, *J* = 7.1 Hz, 1H), 2.78 (s, 6H), 1.63 (d, *J* = 7.1 Hz, 3H). **^13^C NMR**: (150 MHz, CD_3_OD) δ (ppm) = 168.9, 138.8, 136.9, 136.2, 132.2, 132.1, 130.7, 126.6, 126.4, 125.9, 125.5, 117.2, 116.6, 113.2, 111.8, 50.89, 38.56, 17.75 **HRMS**: (ESI) *m/z* calculated for C_19_H_22_N_4_O_3_S [M+H]^+^ 387.1491, found 387.1452

### 3-bromo-N,N-diethylbenzenesulfonamide (29)

The title compound was prepared according to general procedure C, using diethylamine (0.037 mL, 0.36 mmol), dry pyridine (1.8 mL), and 3-bromobenzenesulfonyl chloride (0.056 mL, 0.39 mmol). Workup of the reaction mixture as detailed in general procedure C afforded the aryl-sulfonamide **29** as a colorless oil (97 mg, 92 %). **^1^H NMR**: (500 MHz, CDCl_3_) δ (ppm) = 7.95 (dd, *J* = 1.9, 1.6 Hz, 1H), 7.74 (ddd, *J* = 7.9, 1.6, 1.0 Hz, 1H), 7.67 (ddd, *J* = 8.0, 1.9, 1.0 Hz, 1H), 7.41 (dd, *J* = 8.0, 7.9 Hz, 1H), 3.26 (q, *J* = 7.2 Hz, 4H), 1.15 (t, *J* = 7.2 Hz, 6H).

### N,N-diethyl-3-(4,4,5,5-tetramethyl-1,3,2-dioxaborolan-2-yl)benzenesulfonamide (30)

The title compound was prepared according to general procedure D, using aryl-sulfonamide **29** (141 mg, 0.48 mmol), B_2_Pin_2_ (365.6 mg, 1.44 mmol), NaOAc (188 mg, 1.92 mmol) and Pd(dppf)Cl_2_ (39 mg, 0.05 mmol) in degassed dioxane (1.9 mL). Purification of the crude material by column chromatography (EtOAc:hexanes 10:90) afforded the 3-pinacolboronate aryl-sulfonamide **30** as a colorless solid (94.5 mg, 58 %). **^1^H NMR**: (500 MHz, CDCl_3_) δ (ppm) = 8.23 (dd, *J* = 2.0, 1.0. Hz, 1H), 7.96 (ddd, *J* = 7.4, 1.3, 1.0 Hz, 1H), 7.88 (ddd, *J* = 7.9, 2.0, 1.3 Hz, 1H), 7.48 (dd, *J* = 7.9, 7.4 Hz, 1H), 3.25 (q, *J* = 7.2 Hz, 4H), 1.35 (s, 12H), 1.14 (t, *J* = 7.2 Hz, 6H).

### tert-butyl 5-((tert-butoxycarbonyl)amino)-3-(3-(N,N-diethylsulfamoyl)phenyl)-1H-indole-1-carboxylate (31)

The title compound was prepared according to general procedure A using **7** (128 mg, 0.31 mmol), 3-pinacolboronate aryl-sulfonamide **30** (104 mg, 0.31 mmol), K_2_CO_3_ (127 mg, 0.93 mmol), Pd(PPh_3_)_4_ (35.5 mg, 0.031 mmol), degassed THF (2.6 mL) and degassed water (0.9 mL). Purification by column chromatography (EtOAc:hexanes 10:90) afforded **31** as a pale yellow solid (100 mg, 59%). **^1^H NMR**: (500 MHz, CDCl_3_) δ (ppm) = 8.14 (br t, *J* = 8.0 Hz, 1H), 8.04 (t, *J* = 1.2 Hz, 1H), 7.93 (br s, 1H), 7.80 (tt, *J* = 7.8, 1.2 Hz, 2H), 7.73 (br s, 1H), 7.59 (t, *J* = 7.8 Hz, 1H), 7.23 (dd, *J* = 8.9, 1.8 Hz, 1H), 6.55 (br s, 1H), 3.32 (t, *J* = 7.6 Hz, 4H), 1.69 (s, 9H), 1.52 (s, 9H), 1.19 (t, *J* = 7.6 Hz, 6H).

### (S)-2-amino-N-(3-(3-(N,N-diethylsulfamoyl)phenyl)-1H-indol-5-yl)propanamide (13)

The title compound was prepared according to the general procedures I and B using the Boc-protected-aryl-indole **31** (50 mg, 0.091 mmol) and TFA (0.9 mL) followed by treatment with (*tert*-butoxycarbonyl)*-L*-alanine (20 mg, 0.11 mmol), PyBOP (56 mg, 0.45 mmol), DIPEA (0.08 mL, 0.11 mmol), and dry DMF (0.9 mL). Final Boc-deprotection of the crude mixture using TFA (0.9 mL) and subsequent purification by RP-HPLC (using a SiliCycle SiliaChrom dtC18 semipreparative column (5 μm, 100À, 10 x 250 mm) with a flow rate of 5 mL/min eluting with solvent (A: 0.1 % TFA in water B: 0.1 % TFA in MeCN) on a gradient of (2 → 100) %) solvent B over 15 min, t_R_ = 6.6 min) afforded the TFA salt of **13** as a colorless solid (6 mg, 16%). **^1^H NMR**: (400 MHz, DMSO-d_6_) δ (ppm) = 11.61 (s, 1H), 10.37 (s, 1H), 8.27 (s, 1H), 8.20 (br s, 2H), 7.98 (s, 1H), 7.90 (dt, *J* = 1.6, 7.1 Hz, 1H), 7.88 (d, *J* = 2.5 Hz, 1H), 7.68-7.59 (m, 2H), 7.47 (d, *J* = 8.8 Hz, 1H), 7.31 (dd, *J* = 8.8, 1.6 Hz, 1H), 4.01 (q, *J* = 7.0 Hz, 1H), 3.23 (q, *J* = 7.0 Hz, 4H), 1.47 (d, *J* = 7.0 Hz, 3H), 1.06 (t, *J* = 7.0 Hz, 6H). **^13^C NMR**: (150 MHz, DMSO-d_6_) δ (ppm) = 167.5, 140.4, 136.8, 134.0, 131.1, 130.0, 129.7, 125.5, 124.4, 123.6, 123.1, 115.5, 114.1, 112.3, 109.4, 48.92, 41.83, 17.14, 14.03. **HRMS**: (ESI) *m/z* calculated for C_21_H_27_N_4_O_3_S [M+H]^+^ 415.1804, found 415.

### 1-((3-bromophenyl)sulfonyl)piperidine (32)

The title compound was prepared according to general procedure C, using piperidine (0.035 mL, 0.36 mmol), dry pyridine (1.8 mL), and 3-bromobenzenesulfonyl chloride (0.056 mL, 0.39 mmol). Workup of the reaction mixture as detailed in general procedure C afforded the aryl-sulfonamide **32** as a colorless oil (106 mg, 97 %). **^1^H NMR**: (500 MHz, CDCl_3_) δ (ppm) = 7.90 (dd, *J* = 1.9, 1.7 Hz, 1H), 7.71 (ddd, *J* = 7.9, 1.9, 1.0 Hz, 1H), 7.69 (ddd, *J* = 7.8, 1.7, 1.0 Hz, 1H), 7.41 (dd, *J* = 7.9, 7.8 Hz, 1H), 3.01 (t, *J* = 5.4 Hz, 4H), 1.65 (quint, *J* = 5.4 Hz, 4H), 1.49-1.40 (m, 2H)

### 1-((3-(4,4,5,5-tetramethyl-1,3,2-dioxaborolan-2-yl)phenyl)sulfonyl)piperidine (33)

The title compound was prepared according to general procedure D, using aryl-sulfonamide **32** (208 mg, 0.68 mmol), B_2_Pin_2_ (520 mg, 2.04 mmol), NaOAc (268 mg, 2.72 mmol) and Pd(dppf)Cl_2_ (56 mg, 0.07 mmol) in degassed dioxane (2.7 mL). Purification of the crude material by column chromatography (EtOAc:hexanes 10:90) afforded the 3-pinacolboronate aryl-sulfonamide **33** as a colorless solid (134 mg, 56 %). **^1^H NMR**: (500 MHz, CDCl_3_) δ (ppm) = 8.17 (dd, *J* = 1.8, 1.0. Hz, 1H), 8.00 (ddd, *J* = 7.4, 1.4, 1.0 Hz, 1H), 7.83 (ddd, *J* = 7.9, 1.8, 1.4 Hz, 1H), 7.52 (t, *J* = 7.6 Hz, 1H), 3.00 (t, *J* = 5.4 Hz, 4H), 1.65 (quint, *J* = 5.4 Hz, 4H), 1.45-1.39 (m, 2H), 1.35 (s, 12H).

### tert-butyl 5-((tert-butoxycarbonyl)amino)-3-(3-(piperi-din-1-ylsulfonyl)phenyl)-1H-indole-1-carboxylate (34)

The title compound was prepared according to general procedure A using **7** (117 mg, 0.29 mmol), 3-pinacolboronate aryl-sulfonamide **33** (100 mg, 0.29 mmol), K_2_CO_3_ (118 mg, 0.87 mmol), Pd(PPh_3_)_4_ (33 mg, 0.029 mmol), degassed THF (2.5 mL) and degassed water (0.8 mL). Purification by column chromatography (EtOAc:hexanes 10:90) afforded **34** as a pale yellow solid (110 mg, 68%). **^1^H NMR**: (500 MHz, CDCl_3_) δ (ppm) = 8.13 (br d, *J* = 8.9 Hz, 1H), 8.00 (t, *J* = 1.7 Hz, 1H), 7.98 (br s, 1H), 7.85 (dt, *J* = 7.8, 1.5 Hz, 1H), 7.74 (br s, 1H), 7.72 (dt, *J* = 7.9, 1.7 Hz, 1H), 7.62 (t, *J* = 7.8 Hz, 1H), 7.20 (dd, *J* = 8.9, 2.1 Hz, 1H), 6.56 (br s, 1H), 3.08 (t, *J* = 5.3 Hz, 4H), 1.69 (s, 9H), 1.19 (quint, *J* = 5.3 Hz, 4H), 1.52 (s, 9H), 1.49-1.42 (m, 2H).

### (S)-2-amino-N-(3-(3-(piperidin-1-ylsulfonyl)phenyl)-1H-indol-5-yl)propanamide (14)

The title compound was prepared according to the general procedures I and B using the aryl-indole **34** (49 mg, 0.088 mmol) and TFA (0.9 mL) followed by treatment with (*tert*-butoxycarbonyl)*-L*-alanine (20 mg, 0.11 mmol), PyBOP (55 mg, 0.44 mmol), DIPEA (0.08 mL, 0.11 mmol), and dry DMF (0.9 mL). Final Boc-deprotection of the crude mixture using TFA (0.9 mL) and subsequent purification by RP-HPLC (using a SiliCycle SiliaChrom dtC18 semi-preparative column (5 μm, 100À, 10 x 250 mm) with a flow rate of 5 mL/min eluting with solvent (A: 0.1 % TFA in water B: 0.1 % TFA in MeCN) on a gradient of (2 → 100) %) solvent B over 15 min, t_R_ = 6.7 min) afforded the TFA salt of **14** as a colorless solid (7 mg, 18%). **^1^H NMR**: (400 MHz, CD_3_CN) δ 9.69 (s, 1H), 9.06 (s, 1H), 8.28 (s, 1H), 8.00 (d, *J* = 2.0 Hz, 1H), 7.92 (d, *J* = 7.6 Hz, 1H), 7.70 – 7.58 (m, 3H), 7.48 (d, *J* = 8.7 Hz, 1H), 7.27 (dd, *J* = 8.6, 2.0 Hz, 1H), 4.19 (q, *J* = 7.0 Hz, 1H), 3.03 (t, *J* = 5.7 Hz, 4H), 1.64 (quint, *J* = 5.7 Hz, 4H), 1.59 (d, *J* = 7.0 Hz, 3H), 1.45-1.37 (m, 2H). **^13^C NMR**: (150 MHz, DMSO-d_6_) δ (ppm) = 167.7, 137.0, 136.0, 134.1, 131.4, 130.6, 130.0, 125.9, 124.5, 124.4, 124.0, 115.5, 114.1, 112.5, 109.3, 49.11, 46.90, 24.82, 22.98, 17.39. **HRMS**: (ESI) *m/z* calculated for C_22_H_27_N_4_O_3_S [M+H]^+^ 427.1804, found 427.1753.

### 1-((3-bromophenyl)sulfonyl)pyrrolidine (35)

The title compound was prepared according to general procedure C, using pyrrolidine (0.27 mL, 3.3 mmol), dry pyridine (16.0 mL), and 3-bromobenzenesulfonyl chloride (800 mg, 3.1 mmol). Workup of the reaction mixture as detailed in general procedure C afforded the aryl-sulfonamide **35** as a colorless oil (710 mg, 78 %). **^1^H NMR**: (500 MHz, CDCl_3_) δ (ppm) = 7.97 (dd, *J* = 2.0, 1.7 Hz, 1H), 7.76 (ddd, *J* = 7.8, 1.7, 1.0 Hz, 1H), 7.71 (ddd, *J* = 8.0, 2.0, 1.0 Hz, 1H), 7.41 (dd, *J* = 8.0, 7.8 Hz, 1H), 3.28 – 3.22 (m, 4H), 1.82 – 1.76 (m, 4H). **^13^C NMR**: (125 MHz, CDCl_3_) δ (ppm) = 139.2, 135.7, 130.7, 130.4, 126.1, 123.2, 48.1, 25.4. **HRMS**: (ESI) *m/z* calculated for C_10_H_12_BrNO_2_S [M+NH_4_]^+^ 307.0110, found 307.0112.

### 1-((3-(4,4,5,5-tetramethyl-1,3,2-dioxaborolan-2-yl)phenyl)sulfonyl)pyrrolidine (36)

The title compound was prepared according to general procedure D, using aryl-sulfonamide **35** (686 mg, 2.4 mmol), B_2_pin_2_ (660 mg, 2.60 mmol), NaOAc (485 mg, 5.9 mmol) and Pd(ddpf)Cl_2_ (97 mg, 0.12 mmol) in degassed DMSO (12 mL). Purification of the crude material by column chromatography (EtOAc:hexanes 16:84) afforded **36** as a colorless solid (493 mg, 65 %). **^1^H NMR**: (500 MHz, CDCl_3_) δ (ppm) = 8.25 (dd, *J* = 2.0, 1.4 Hz, 1H), 7.99 (ddd, *J* = 7.4, 1.4, 1.3 Hz, 1H), 7.90 (ddd, *J* = 7.9, 2.0, 1.3 Hz, 1H), 7.51 (dd, *J* = 7.9, 7.4 Hz, 1H), 3.29 – 3.22 (m, 4H), 1.78 – 1.70 (m, 4H), 1.34 (s, 12H). **^13^C NMR**: (125 MHz, CDCl_3_) δ (ppm) = 138.8, 136.6, 133.7, 130.1, 128.4, 84.5, 48.1, 25.3, 25.0. *13C-B(OR)2 not ob-served. **HRMS**: (ESI) *m/z* calculated for C_16_H_24_BNO_4_S [M+H]^+^ 338.1529, found 338.1608.

### 3-(3-(pyrrolidin-1-ylsulfonyl)phenyl)-1H-indol-5-amine (37)

The title compound was prepared according to general procedure A using **7** (100 mg, 0.24 mmol), 3-pinacolboronate aryl-sulfonamide **36** (82 mg, 0.24 mmol), K_2_CO_3_ (101 mg, 0.75 mmol), Pd(PPh_3_)_4_ (28 mg, 0.024 mmol), degassed THF (2.07 mL) and degassed water (0.7 mL). Purification by column chromatography afforded the protected coupled product, which was subsequently deprotected in TFA (2.4 mL) and concentrated under reduced pressure to afford the TFA salt of **37** as a brown solid (40 mg, 36%). **^1^H NMR**: (500 MHz, CD_3_OD) δ (ppm) = 8.09 (s, 1H), 7.95 (d, *J* = 7.6 Hz, 1H), 7.84 (s, 1H), 7.79 (s, 1H), 7.74 (d, *J* = 7.8 Hz, 1H), 7.69 (dd, *J* = 7.8, 7.6 Hz, 1H), 7.61 (d, *J* = 8.6 Hz, 1H), 7.18 (dd, *J* = 8.6, 2.0 Hz, 1H), 3.32 – 3.28 (s, 4H), 1.80 – 1.76 (m, 4H). **^13^C NMR**: (125 MHz, CD3OD) δ (ppm) = 138.7, 138.1, 137.9, 132.3, 131.0, 127.3, 126.8, 126.4, 125.7, 125.6, 117.3, 116.8, 114.5, 113.5, 49.3, 26.3. **HRMS**: (ESI) *m/z* calculated for C_18_H_19_N_3_O_2_S [M+H]^+^ 342.1271, found 342.1276.

### (S)-2-amino-N-(3-(3-(pyrrolidin-1-ylsulfonyl)phenyl)-1H-indol-5-yl)propanamide (15)

The title compound was prepared according to general procedure B using the aryl-indole **37** (27 mg, 0.06 mmol), (*tert*-butoxycarbonyl)*-L*-alanine (11 mg, 0.06 mmol), PyBOP (31 mg, 0.06 mmol), DIPEA (0.08 mL, 0.30 mmol), and dry DMF (0.6 mL). RP-HPLC (gradient: 2-50 shortprep, tR = 8.99 min) of the crude deprotected product afforded the TFA salt of **15** as a colorless solid (11 mg, 34%). **^1^H NMR**: (500 MHz, CD_3_OD) δ (ppm) = 8.29 (s, 1H), 8.10 (s, 1H), 7.94 (d, *J* = 7.6 Hz, 1H), 7.68 (d, *J* = 7.7 Hz, 1H), 7.644 (s, 1H), 7.636 (dd, *J* = 7.7, 7.6 Hz, 1H), 7.45 (d, *J* = 8.7 Hz, 1H), 7.27 (dd, *J* = 8.7 Hz, 1H), 4.09 (q, *J* = 7.0 Hz, 1H), 3.34 – 3.30 (m, 4H), 1.79 – 1.75 (m, 4H), 1.63 (d, *J* = 7.0 Hz, 3H). **^13^C NMR**: (125 MHz, CD_3_OD) δ (ppm) = 169.0, 138.8, 138.3, 136.2, 132.1, 132.0, 130.7, 126.4, 126.3, 125.9, 125.3, 117.3, 116.6, 113.2, 111.9, 50.9, 49.4, 26.2, 17.8. **HRMS**: (ESI) *m/z* calculated for C_21_H_2_4N4O_3_S [M+H]^+^ 413.1642, found 413.1643.

### 1-bromo-3-(isopropylsulfonyl)benzene (38)

The title compound was prepared according to general procedure E, using 3-bromothiophenol (0.66 mL, 6.4 mmol, 1.0 equiv.), K_2_CO_3_ (1.23 g, 8.89 mmol, 1.4 equiv.), 2-bromopropane (0.72 mL, 7.6 mmol, 1.2 equiv.) in dry acetone (21 mL, 0.3 M). Oxidation was then performed according to general procedure E using MeOH (36 mL, 0.17 M according to the sulfide), oxone (5.593 g, 18 mmol, 3.0 equiv.), and water (36 mL, 0.5 M according to the oxone). Work-up and concentration of the crude reaction mixture afforded **38** as a yellow oil (1.10 g, 69 % over 2 steps). **^1^H NMR**: (500 MHz, CDCl_3_) δ (ppm) = 8.00 (dd, *J* = 1.9, 1.7 Hz, 1H), 7.79 (ddd, *J* = 7.9, 1.7, 1.1 Hz, 1H), 7.76 (ddd, *J* = 8.0, 2.0, 1.0 Hz, 1H), 7.43 (dd, *J* = 8.0, 7.9 Hz, 1H), 3.19 (septet, *J* = 6.9 Hz, 1H), 1.28 (d, *J* = 6.9 Hz, 6H). **^13^C NMR**: (125 MHz, CDCl_3_) δ (ppm) = 139.1, 136.8, 131.9, 130.7, 127.7, 123.2, 55.8, 15.7.

### 2-(3-(isopropylsulfonyl)phenyl)-4,4,5,5-tetramethyl-1,3,2-dioxaborolane (39)

The title compound was prepared according to general procedure D, using isopropyl sulfone **38** (1.03 g, 3.92 mmol), B_2_pin_2_ (1.10 g, 4.32 mmol), NaOAc (0.805 g, 9.81 mmol) and Pd(ddpf)Cl_2_ (0.160 g, 0.20 mmol) in degassed DMSO (20 mL). Purification of the crude material by column chromatography (EtOAc:hexanes 1:4) afforded the 3-pinacolboronate aryl-sulfone **39** as a colorless solid (0.805 g, 66 %). **^1^H NMR**: (500 MHz, CDCl_3_) δ (ppm) = 8.30 (dd, *J* = 2.0, 1.3 Hz, 1H), 8.05 (ddd, *J* = 7.3, 1.3, 1.3 Hz, 1H), 7.95 (ddd, *J* = 7.9, 2.0, 1.3 Hz, 1H), 7.55 (dd, *J* = 7.9, 7.3 Hz, 1H), 3.21 (septet, *J* = 6.9 Hz, 1H), 1.34 (s, 12H), 1.29 (d, *J* = 6.9 Hz, 6H). **^13^C NMR**: (125 MHz, CDCl_3_) δ (ppm) = 139.8, 136.8, 135.1, 131.6, 128.4, 84.6, 55.5, 25.0, 15.8. *13C-B(OR)2 not observed. **HRMS**: (ESI) *m/z* calculated for C_15_H_23_BO_4_S [M+NH_4_]^1^ 328.1748, found 328.1760.

### 3-(3-(isopropylsulfonyl)phenyl)-1H-indol-5-amine (40)

The title compound was prepared according to general procedure A using **7** (597 mg, 1.45 mmol), isopropyl sulfone **39** (450 mg, 1.45 mmol), K_2_CO_3_ (602 mg, 4.35 mmol), Pd(PPh_3_)_4_ (168 mg, 0.145 mmol), degassed THF (12 mL) and degassed water (4 mL). Purification by column chromatography afforded the protected coupled product, which was subsequently deprotected in TFA (15 mL) and concentrated under reduced pressure to afford the TFA salt of **40** as a brown solid (479 mg, 77%). **^1^H NMR**: (500 MHz, CD_3_OD) δ (ppm) = 8.13 (dd, *J* = 1.8, 1.8 Hz, 1H), 8.02 (ddd, *J* = 7.7, 1.8, 1.4 Hz, 1H), 7.89 (d, *J* = 2.1 Hz, 1H), 7.80 (s, 1H), 7.78 (ddd, *J* = 7.9, 1.8, 1.4 Hz, 1H), 7.71 (dd, *J* = 7.9, 7.7 Hz, 1H), 7.63 (d, *J* = 8.6 Hz, 1H), 7.22 (dd, *J* = 8.6, 2.1 Hz, 1H), 3.40 (septet, *J* = 6.8 Hz, 1H), 1.30 (d, *J* = 6.8 Hz, 6H). **^13^C NMR**: (125 MHz, CD_3_OD) δ (ppm) = 139.0, 138.18, 138.16, 133.2, 131.0, 127.7, 127.5, 127.1, 126.7, 124.6, 117.5, 116.6, 114.6, 114.1, 56.5, 15.9. **HRMS**: (ESI) *m/z* calculated for C_17_H_18_N_2_O_2_S [M+H]^+^ 315.1162, found 315.1170.

### (S)-2-amino-N-(3-(3-(isopropylsulfonyl)phenyl)-1H-in-dol-5-yl)propanamide (16)

The title compound was prepared according to general procedure B using the aryl-indole **40** (56 mg, 0.14 mmol), (*tert*-butoxycarbonyl)-*L*-alanine (30 mg, 0.16 mmol), PyBOP (85 mg, 0.16 mmol), DIPEA (0.18 mL, 0.68 mmol), and dry DMF (1.4 mL). Purification by RP-HPLC (using a SiliCycle SiliaChrom dtC18 semipreparative column (5 μm, 100Å, 10 x 250 mm) with a flow rate of 5 mL/min eluting with solvent (A: 0.1 % TFA in water B: 0.1 % TFA in MeCN) on a gradient of (2 → 100) % solvent B over 15 min, tR = 6.11 min) of the crude deprotected product afforded the TFA salt of **16** as a colorless solid (10 mg, 15%). **^1^H NMR**: (600 MHz, CD_3_CN) δ (ppm) = 9.71 (s, 1H), 9.31 (s, 1H), 8.19 (d, *J* = 2.0 Hz, 1H), 8.07 (t, *J* = 1.8 Hz, 1H), 7.95 (ddd, *J* = 7.7, 1.5, 1.4 Hz, 1H), 7.71 (ddd, *J* = 7.9, 1.5, 1.4 Hz, 1H), 7.65 (dd, *J* = 7.9, 7.7 Hz, 1H), 7.63 (d, *J* = 2.7 Hz, 1H), 7.44 (d, *J* = 8.7 Hz, 1H), 7.33 (dd, *J* = 8.7, 2.0 Hz, 1H), 4.24 (q, *J* = 7.1 Hz, 1H), 3.35 (septet, *J* = 6.8 Hz, 1H), 1.60 (d, *J* = 7.1 Hz, 3H), 1.26 (d, *J* = 6.8 Hz, 6H). **^13^C NMR**: (150 MHz, CD_3_CN) δ (ppm) = 168.5, 138.7, 137.9, 135.2, 132.6, 132.3, 130.6, 127.4, 126.6, 125.9, 125.8, 116.9, 116.0, 113.2, 110.9, 55.9, 50.9, 17.6, 15.88, 15.87. **HRMS**: (ESI) *m/z* calculated for C_20_H_23_N_3_O_3_S [M+H]^+^ 386.1533, found 386.1538.

### 1-bromo-3-(cyclopentylsulfonyl)benzene (41)

The title compound was prepared according to general procedure E, using 3-bromothiophenol (0.44 mL, 4.2 mmol, 1.0 equiv.), Cs2CO_3_ (2.99 g, 8.5 mmol, 2.0 equiv.), bromocyclopentane (0.43 mL, 4.2 mmol, 1.0 equiv.) in dry DMF (21 mL, 0.2 M). Oxidation of the crude sulfide (*ca*. 713 mg, 2.8 mmol, 1.0 equiv.) was then performed according to general procedure E using MeOH (17.0 mL, 0.165 according to the sulfide), and oxone (2.56 g, 8.3 mmol, 3.0 equiv.) in water (17 mL, 0.5 M). Work-up and concentration of the crude reaction mixture afforded the cyclopentyl sulfone **41** as a colorless oil (600 mg, 49 % over 2 steps). **^1^H NMR**: (500 MHz, CDCl_3_) δ (ppm) = 8.02 (dd, *J* = 2.0, 1.8 Hz, 1H), 7.81 (ddd, *J* = 7.8, 1.8, 1.1 Hz, 1H), 7.74 (ddd, *J* = 8.0, 2.0, 1.0 Hz, 1H), 7.42 (dd, *J* = 8.0, 7.8 Hz, 1H), 3.47 (tt, *J* = 8.7, 7.1 Hz, 1H), 2.11 – 1.96 (m, 2H), 1.92 – 1.81 (m, 2H), 1.80 – 1.70 (m, 2H), 1.66 – 1.53 (m, 2H). **^13^C NMR**: (125 MHz, CDCl_3_) δ (ppm) = 141.2, 136.6, 131.4, 130.8, 127.1, 123.3, 64.4, 27.3, 25.9. **HRMS**: (ESI) *m/z* calculated for C11H13BrO_2_S [M+NH_4_]^+^ 306.0158, found 306.0171.

### 2-(3-(cyclopentylsulfonyl)phenyl)-4,4,5,5-tetramethyl-1,3,2-dioxaborolane (42)

The title compound was prepared according to general procedure D, using cyclopentyl sulfone **41** (520 mg, 1.8 mmol), B_2_pin_2_ (502 mg, 1.98 mmol), NaOAc (369 mg, 4.5 mmol) and Pd(ddpf)Cl_2_ (73 mg, 0.090 mmol) in degassed DMSO (9.0 mL). Purification of the crude material by column chromatography (EtOAc:hexanes 1:4) afforded the 3-pinacolboronate aryl-sulfone **42** as a yellow solid (438 mg, 72 %). **^1^H NMR**: (500 MHz, CDCl_3_) δ (ppm) = 8.32 (dd, *J* = 2.0, 1.4 Hz, 1H), 8.03 (ddd, *J* = 7.4, 1.4, 1.2 Hz, 1H), 7.97 (ddd, *J* = 7.9, 2.0, 1.2 Hz, 1H), 7.54 (dd, *J* = 7.9, 7.4 Hz, 1H), 3.51 (tt, *J* = 8.8, 7.2 Hz, 1H), 2.11 – 2.03 (m, 2H), 1.90 – 1.81 (m, 2H), 1.81 – 1.73 (m, 2H), 1.65 – 1.54 (m, 2H), 1.34 (s, 12H). **^13^C NMR**: (125 MHz, CDCl_3_) δ (ppm) = 139.7, 138.7, 134.6, 131.0, 128.5, 84.5, 64.2, 27.3, 26.0, 25.0. *13C-B(OR)2 not observed. **HRMS**: (ESI) *m/z* calculated for C_17_H_25_BO_4_S [M+NH_4_p 354.1905, found 354.1933.

### 3-(3-(cyclopentylsulfonyl)phenyl)-1H-indol-5-amine (43)

The title compound was prepared according to general procedure A using **7** (200 mg, 0.49 mmol), cyclopentyl sulfone **42** (164 mg, 0.49 mmol), K_2_CO_3_ (202 mg, 1.46 mmol), Pd(PPh_3_)_4_ (56 mg, 0.049 mmol), degassed THF (4.14 mL) and degassed water (1.4 mL). Purification by column chromatography afforded the protected coupled product, which was subsequently deprotected in TFA (4.8 mL) and concentrated under reduced pressure to afford the TFA salt of **43** as a yellow solid (141 mg, 64%). **^1^H NMR**: (500 MHz, CD_3_OD) δ (ppm) = 8.16 (s, 1H), 8.00 (d, *J* = 7.8 Hz, 1H), 7.89 (d, *J* = 2.1 Hz, 1H), 7.90 – 7.88 (m, 2H), 7.71 (dd, *J* = 7.8, 7.6 Hz, 1H), 7.63 (d, *J* = 8.6 Hz, 1H), 7.22 (dd, *J* = 8.6, 2.1 Hz, 1H), 3.76 (ddd, *J* = 15.8, 8.9, 6.9 Hz, 1H), 2.11 – 1.99 (m, 2H), 1.96 – 1.86 (m, 2H), 1.81 – 1.71 (m, 2H), 1.69 – 1.61 (m, 2H). **NMR**: (125 MHz, CD_3_OD) δ (ppm) = 140.9, 138.23, 138.19, 133.1, 131.1, 127.5, 127.1, 126.8, 126.6, 124.6, 117.5, 116.7, 114.6, 114.1, 65.0, 28.2, 26.9. **HRMS**: (ESI) *m/z* calculated for C_19_H_20_N_2_O_2_S [M+H]^+^ 341.1318, found 341.1318.

## Synthesis of sulfone 17

The title compound was prepared according to general procedure B using the aryl-indole **43** (45 mg, 0.10 mmol), (*tert*-butoxycarbonyl) *-L*-alanine (19 mg, 0.10 mmol), PyBOP (52 mg, 0.10 mmol), DIPEA (0.14 mL, 0.50 mmol), and dry DMF (1.0 mL). RP-HPLC (gradient: 2-50 shortprep, tR = 7.52 min) of the crude deprotected product afforded the TFA salt of **17** as a colorless solid (27 mg, 50%). **^1^H NMR**: (500 MHz, CD_3_CN) δ (ppm) = 9.74 (s, 1H), 9.18 (s, 1H), 8.17 (d, *J* = 1.9 Hz, 1H), 8.09 (dd, *J* = 1.8, 1.4 Hz, 1H), 7.93 (dd, *J* = 7.8, 1.4, 1.4 Hz, 1H), 7.72 (dd, *J* = 7.9, 1.4, 1.4 Hz, 1H), 7.64 (dd, *J* = 7.9, 7.8 Hz, 1H), 7.62 (d, *J* = 2.6 Hz, 2H), 7.45 (d, *J* = 8.7 Hz, 1H), 7.30 (dd, *J* = 8.8, 2.4 Hz, 1H), 4.23 (q, *J* = 7.0 Hz, 1H), 3.68 (tt, *J* = 8.9, 7.0 Hz, 1H), 2.04 – 1.96 (m, 2H), 1.91 – 1.82 (m, 2H), 1.73 – 1.66 (m, 2H), 1.63 – 1.55 (m, 5H). **^13^C NMR**: (125 MHz, CD_3_CN) δ (ppm) = 168.6, 140.7, 138.1, 135.3, 132.5, 132.2, 130.7, 126.9, 126.1, 126.0, 125.9, 117.1, 116.1, 113.2, 111.2, 64.6, 51.0, 28.0, 27.9, 26.6, 17.6. **HRMS**: (ESI) *m/z* calculated for C_22_H_25_N_3_O_3_S [M+H]^+^ 412.1689, found 412.1706.

### (S)-2-amino-N-(3-(3-(cyclopentylsulfonyl)phenyl)-1H-indol-5-yl)propanamide (18a)

The title compound was prepared according to general procedure B using the aryl-indole **24** (100 mg, 0.25 mmol), (*tert*-butoxycarbonyl)-*L*-proline (65 mg, 0.30 mmol), PyBOP (156 mg, 0.30 mmol), DIPEA (0.33 mL, 1.25 mmol), and dry DMF (2.5 mL). Purification by RP-HPLC (using a SiliCycle SiliaChrom dtC18 semipreparative column (5 μm, 100À, 10 x 250 mm) with a flow rate of 5 mL/min eluting with solvent (A: 0.1 % TFA in water B: 0.1 % TFA in MeCN) on a gradient of (2 → 100) % solvent B over 15 min, tR = 5.88 min) of the crude deprotected product afforded the TFA salt of **18a** as a pale yellow solid (21 mg, 17%). **^1^H NMR**: (600 MHz, CD_3_OD) δ (ppm) = 8.25 (d, *J* = 1.9 Hz, 1H), 8.22 (dd, *J* = 1.9, 1.9 Hz, 1H), 7.99 (ddd, *J* = 7.7, 1.9, 1.8 Hz, 1H), 7.80 (ddd, *J* = 7.6, 1.9, 1.8 Hz, 1H), 7.67 (s, 1H), 7.65 (dd, *J* = 7.7, 7.6 Hz, 1H), 7.45 (d, *J* = 8.7 Hz, 1H), 7.34 (dd, *J* = 8.7, 1.9 Hz, 1H), 4.43 (dd, *J* = 7.7, 7.7 Hz, 1H), 3.48 (ddd, *J* = 11.4, 7.1, 7.0 Hz, 1H), 3.39 (ddd, *J* = 11.4, 71, 7.0 Hz, 1H), 3.19 (s, 3H), 2.55 (dddd, *J* = 13.6, 7.1, 7.0, 7.0 Hz, 1H), 2.26 – 2.04 (m, 3H). **^13^C NMR**: (150 MHz, CD_3_OD) δ (ppm) = 167.7, 142.5, 139.1, 136.2, 132.9, 132.1, 131.0, 126.3, 126.0, 126.0, 124.9, 117.2, 116.4, 113.2, 111.8, 61.7, 47.5, 44.5, 31.2, 25.2. **HRMS**: (ESI) *m/z* calculated for C_20_H_21_N_3_O_3_S [M+H]^+^ 384.1376, found 384.1386.

### N1-(3-(3-(methylsulfonyl)phenyl)-1H-indol-5-yl)ethane-1,2-diamine (19a)

To a stirred solution of the aryl-indole **24** (35 mg, 0.12 mmol, 2.0 equiv.) in water (*ca*. 0.15 mL, 2.0 M) was added 2-bromoethan-1-amine hydrochloride (13 mg, 0.061 mmol, 1.0 equiv.) at room temperature. The reaction mixture was then stirred at 95 °C for 22 hours and cooled to room temperature. The reaction mixture was then diluted with water and extracted with EtOAc (3 times). The remaining aqueous layer was purified by RP-HPLC (gradient: 2-30 shortprep, tR = 8.33 min) to afford the TFA salt of **19a** as a brown gum (4 mg, 13%). **^1^H NMR**: (500 MHz, CD_3_CN) δ (ppm) = 9.47 (s, 1H), 8.17 (dd, *J* = 1.9, 1.4 Hz, 1H), 8.01 (ddd, *J* = 7.8, 1.4, 1.1 Hz, 1H), 7.76 (ddd, *J* = 7.8, 1.9, 1.1 Hz, 1H), 7.67 (dd, *J* = 7.8, 7.8 Hz, 1H), 7.57 (d, *J* = 2.6 Hz, 1H), 7.36 (d, *J* = 8.5 Hz, 1H), 7.16 (s, 1H), 6.75 (d, *J* = 8.5 Hz, 1H), 3.49 (t, *J* = 5.9 Hz, 2H), 3.22 (t, *J* = 5.9 Hz, 2H), 3.12 (s, 3H).

### N1-methyl-N2-(3-(3-(methylsulfonyl)phenyl)-1H-indol-5-yl)ethane-1,2-diamine (19b)

To a stirred solution of the arylindole **24** (22 mg, 0.077 mmol, 1.0 equiv.) in 1,2-dichloroethane (0.77 mL, 0.1 M) was added *tert*-butyl methyl(2-oxoethyl)carbamate (13 mg, 0.077 mmol, 1.0 equiv.) at room temperature. The reaction mixture was then stirred at room temperature for 30 minutes, after which NaBH(OAc)_3_ (24 mg, 0.12 mmol, 1.5 equiv.) was added. The solution was then stirred for 18 hours, diluted with saturated aqueous NaHCO_3_ and extracted with CH_2_Cl_2_ (3 times). The combined organic layers were dried over MgSO_4_ and concentrated under reduced pressure to afford the crude protected indole (brown gum) which was used in the next step without further purification. The crude protected product was dissolved in TFA (neat, 0.77 mL, 0.1 M) and stirred at room temperature for 30 minutes. The reaction mixture was then concentrated under reduced pressure to afford the crude product, which was purified by RP-HPLC (using a SiliCycle SiliaChrom dtC18 semipreparative column (5 μm, 100À, 10 x 250 mm) with a flow rate of 5 mL/min eluting with solvent (A: 0.1 % TFA in water B: 0.1 % TFA in MeCN) on a gradient of (2 → 30) % solvent B over 15 min, tR = 8.95 min) to afford the TFA salt of **19b** as a brown gum (3 mg, 8%). **^1^H NMR**: (600 MHz, CD_3_OD) δ (ppm) = 8.26 (s, 1H), 7.99 (d, *J* = 8.0 Hz, 1H), 7.79 (d, *J* = 7.8 Hz, 1H), 7.68 (dd, *J* = 7.8, 7.8 Hz, 1H), 7.58 (s, 1H), 7.34 (d, *J* = 8.7 Hz, 1H), 7.23 (d, *J* = 2.1 Hz, 0H), 6.80 (dd, *J* = 8.7, 2.1 Hz, 1H), 3.53 (t, *J* = 6.0 Hz, 2H), 3.31 (t, *J* = 6.1 Hz, 2H), 3.19 (s, 3H), 2.78 (s, 3H). **^13^C NMR**: (150 MHz, CD_3_OD) δ (ppm) = 143.0, 142.4, 139.7, 133.6, 132.6, 131.0, 127.1, 125.8, 125.1, 124.5, 115.4, 113.9, 113.8, 102.5, 49.6, 44.4, 42.9, 33.7. **HRMS**: (ESI) *m/z* calculated for C_18_H_21_N_3_O_2_S [M+H]^+^ 344.1427, found 344.1427.

### (S)-N-(3-(3-(isopropylsulfonyl)phenyl)-1H-indol-5-yl)-2-(methylamino)propanamide (18b)

The title compound was prepared according to general procedure B using the aryl-indole **40** (50 mg, 0.12 mmol), *N*-(*tert*-butoxycarbonyl)-*N*-methyl*-L*-alanine (25 mg, 0.12 mmol), PyBOP (63 mg, 0.12 mmol), DIPEA (0.16 mL, 0.61 mmol), and dry DMF (1.22 mL). RP-HPLC (gradient: 2-30 shortprep, tR = 11.84 min) of the crude deprotected product afforded the TFA salt of **18b** as a colorless solid (19 mg, 30%). **^1^H NMR**: (600 MHz, CD_3_CN) δ (ppm) = 9.86 (s, 1H), 9.39 (s, 1H), 8.19 (s, 1H), 8.07 (d, *J* = 1.8 Hz, 1H), 7.95 (dd, *J* = 7.6, 1.7 Hz, 1H), 7.70 (dd, *J* = 7.8, 1.7 Hz, 1H), 7.65 (dd, *J* = 7.8, 7.6 Hz, 1H), 7.62 (d, *J* = 2.5 Hz, 1H), 7.46 (d, *J* = 8.6 Hz, 1H), 7.32 (d, *J* = 8.7 Hz, 1H), 4.06 (q, *J* = 7.0 Hz, 1H), 3.33 (*septet, J* = 6.7 Hz, 1H), 2.68 (s, 3H), 1.59 (d, *J* = 6.8 Hz, 3H), 1.25 (d, *J* = 6.8 Hz, 6H). **^13^C NMR**: (150 MHz, CD3CN) δ (ppm) = 167.8, 138.7, 137.9, 135.3, 132.7, 132.0, 130.6, 127.4, 126.6, 126.0, 125.8, 117.1, 116.0, 113.2, 111.3, 58.8, 56.0, 32.1, 16.3, 15.91, 15.90. **HRMS**: (ESI) *m/z* calculated for C_21_H_2_5N3O_3_S [M+H]^+^ 400.1689, found 400.1703.

### (S)-N-(3-(3-(isopropylsulfonyl)phenyl)-1H-indol-5-yl)azetidine-2-carboxamide (18c)

The title compound was prepared according to general procedure B using the aryl-indole **40** (50 mg, 0.122 mmol), *N*-Boc*-L*-azetidine-2-carboxylic acid (25 mg, 0.122 mmol), PyBOP (63 mg, 0.122 mmol), DIPEA (0.16 mL, 0.61 mmol), and dry DMF (1.22 mL). Purification by RP-HPLC (using a SiliCycle SiliaChrom dtC18 semipreparative column (5 μm, 100Å, 10 x 250 mm) with a flow rate of 5 mL/min eluting with solvent (A: 0.1 % TFA in water B: 0.1 % TFA in MeCN) on a gradient of (2 → 30) % solvent B over 15 min, tR = 11.69 min) of the crude deprotected product afforded the TFA salt of **18c** as a colorless solid (14 mg, 23%). **^1^H NMR**: (600 MHz, CD_3_CN) δ (ppm) = 9.81 (s, 1H), 9.36 (s, 1H), 8.21 (s, 1H), 8.08 (s, 1H), 7.96 (d, *J* = 7.5 Hz, 1H), 7.72 (d, *J* = 7.7 Hz, 1H), 7.66 (dd, *J* = 7.7, 7.5 Hz, 1H), 7.63 (s, 1H), 7.47 (d, *J* = 8.6 Hz, 1H), 7.32 (d, *J* = 8.7 Hz, 1H), 5.23 (dd, *J* = 9.4, 7.7 Hz, 1H), 4.16 (q, *J* = 9.3 Hz, 1H), 3.96 (td, *J* = 10.1, 6.4 Hz, 1H), 3.35 (*septet, J* = 6.4 Hz, 1H), 2.82 (qd, *J* = 10.1, 6.5 Hz, 1H), 2.64 (dt, *J* = 18.6, 8.4 Hz, 1H), 1.27 (d, *J* = 6.5 Hz, 6H). **^13^C NMR**: (150 MHz, CD3CN) δ (ppm) = 166.4, 138.8, 138.0, 135.3, 132.7, 132.2, 130.6, 127.4, 126.6, 126.0, 125.9, 116.9, 116.1, 113.3, 111.0, 59.6, 56.0, 44.7, 24.0, 15.9. **HRMS**: (ESI) *m/z* calculated for C_21_H_23_N_3_O_3_S [M+H]^+^ 398.1533, found 398.1564.

### 3-(5-amino-1H-indol-3-yl)-N,N-dimethylbenzenesulfonamide (44)

The title compound was prepared according to general procedure A using **7** (400 mg, 0.97 mmol), N,N-dimethyl-3-(4,4,5,5-tetramethyl-1,3,2-dioxaborolan-2-yl)benzene-sulfonamide (303 mg, 0.97 mmol), K_2_CO_3_ (403 mg, 2.9 mmol), Pd(PPh_3_)_4_ (112 mg, 0.097 mmol), degassed THF (8.3 mL) and degassed water (2.8 mL). Purification by column chromatography afforded the protected coupled product, which was subsequently deprotected in TFA (10 mL) and concentrated under reduced pressure to afford the TFA salt of **44** as a brown solid (261 mg, 62%). **^1^H NMR**: (500 MHz, CD3OD) δ (ppm) = 8.03 (s, 1H), 7.96 (ddd, *J* = 7.2, 1.8 1.7 Hz, 1H), 7.88 (d, *J* = 2.1 Hz, 1H), 7.79 (d, *J* = 1.5 Hz, 1H), 7.72 – 7.66 (m, 2H), 7.63 (d, *J* = 8.6 Hz, 1H), 7.22 (dd, *J* = 8.7, 2.1 Hz, 1H), 2.75 (s, 6H). **^13^C NMR**: (125 MHz, CD_3_OD) δ (ppm) = 138.2, 138.0, 137.2, 132.4, 130.9, 127.4, 126.8, 126.7, 126.0, 124.5, 117.5, 116.8, 114.6, 114.0, 38.4. **HRMS**: (ESI) *m/z* calculated for C16H17N3O_2_S [M+H]^+^ 316.1114, found 316.1120.

### (S)-N-(3-(3-(N,N-dimethylsulfamoyl)phenyl)-1H-indol-5-yl)azetidine-2-carboxamide (18d)

The title compound was prepared according to general procedure B using the aryl indole **44** (58mg, 0.14mmol), *N*-Boc-*L*-azetidine-2-carboxylic acid (27mg, 0.14mmol), PyBOP (70mg, 0.14mmol), DIPEA (0.179mL, 0.68mmol) and dry DMF (1.45mL). Purification by RP-HPLC (using a SiliCycle SiliaChrom dtC18 semipreparative column (5 μm, 100Å, 10 x 250 mm) with a flow rate of 5 mL/min eluting with solvent (A: 0.1 % TFA in water B: 0.1 % TFA in MeCN) on a gradient of (2 → 50) % solvent B over 15 min, tR = 8.60 min) of the crude deprotected product afforded the TFA salt of **18d** as a colorless solid (14 mg, 23%) (14mg, 20%). **^1^H NMR:** (400MHz, CD_3_CN) δ (ppm) = δ 9.77 (s, 1H), 9.06 (s, 1H), 8.27 (d, *J* = 1.8 Hz, 1H), 8.01 (t, *J* = 1.7, 1H), 7.96 – 7.86 (m, 1H), 7.70 – 7.59 (m, 3H), 7.44 (d, *J* = 8.7 Hz, 1H), 7.26 (dd, *J* = 8.7, 2.0 Hz, 1H), 5.16 (t, *J* = 8.6 Hz, 1H), 4.31 – 3.82 (m, 2H), 2.91 – 2.76 (m, 1H), 2.74 (s, 6H), 2.72 – 2.55 (m, 1H). **^13^C NMR**: (150 MHz, CD_3_CN) δ (ppm) = 137.9, 136.6, 135.2, 132.3, 132.0, 130.7, 126.4, 126.0, 125.7, 116.7, 113.3, 110.7, 59.8, 44.7, 41.3, 38.7, 24.6. *^13^C=O not observed. **HRMS:** (ESI) *m/z* calculated for C_20_H_2_3N4O_3_S [M+H]^+^ 399.1485, found 399.1490.

### (S)-N-(3-(3-(N,N-dimethylsulfamoyl)phenyl)-1H-indol-5-yl)-2-(methylamino)propanamide (18e)

The title compound was prepared according to general procedure B using the aryl-indole **44** (50 mg, 0.116 mmol), *N*-(*tert*-butoxycarbonyl)-*N*-methyl*-L*-alanine (24 mg, 0.116 mmol), PyBOP (61 mg, 0.116 mmol), DIPEA (0.15 mL, 0.58 mmol), and dry DMF (1.2 mL). Purification by RP-HPLC (using a SiliCycle SiliaChrom dtC18 semipreparative column (5 μm, 100Å, 10 x 250 mm) with a flow rate of 5 mL/min eluting with solvent (A: 0.1 % TFA in water B: 0.1 % TFA in MeCN) on a gradient of (2 → 30) % solvent B over 15 min, tR = 11.84 min) of the crude deprotected product afforded the TFA salt of **18e** as a colorless solid (20 mg, 34%). **^1^H NMR**: (500 MHz, CD_3_OD) δ (ppm) = 8.31 (d, *J* = 2.0 Hz, 1H), 8.06 (dd, *J* = 1.8 Hz, 1H), 7.96 (ddd, *J* = 7.2, 1.8, 1.7 Hz, 1H), 7.70 – 7.61 (m, 3H), 7.45 (d, *J* = 8.7 Hz, 1H), 7.27 (dd, *J* = 8.7, 2.0 Hz, 1H), 3.87 (q, *J* = 7.0 Hz, 1H), 2.77 (s, 6H), 2.70 (s, 3H), 1.60 (d, *J* = 7.0 Hz, 3H). **^13^C NMR**: (150 MHz, CD3OD) δ (ppm) = 169.14, 138.8, 136.8, 136.2, 132.2, 132.0, 130.7, 126.6, 126.4, 125.9, 125.6, 117.1, 116.5, 113.2, 111.8, 59.3, 38.6, 32.2, 16.9. **HRMS**: (ESI) *m/z* calculated for C_20_H_24_N_4_O_3_S [M+H]^+^ 401.1642, found 401.1655.

### 2-amino-N-(3-(3-(N,N-dimethylsulfamoyl)phenyl)-1H-indol-5-yl)acetamide (18f)

The title compound was prepared according to general procedure B using the aryl-indole **44** (45 mg, 0.10 mmol), (*tert*-butoxycarbonyl)glycine (18 mg, 0.10 mmol), PyBOP (54 mg, 0.10 mmol), DIPEA (0.14 mL, 0.52 mmol), and dry DMF (1.0 mL). RP-HPLC (gradient: 2-50 shortprep, tR = 10.38 min) of the crude deprotected product afforded the TFA salt of **18f** as a colorless solid (13 mg, 25%). **^1^H NMR**: (500 MHz, CD_3_OD:CD_3_CN 1:1, calibrated to CD_3_OD) δ (ppm) = 8.32 (d, *J* = 2.0 Hz, 1H), 8.07 (s, 1H), 7.98 (d, *J* = 7.2 Hz, 1H), 7.72 – 7.65 (m, 3H), 7.50 (d, *J* = 8.7 Hz, 1H), 7.29 (dd, *J* = 8.7, 2.0 Hz, 1H), 3.85 (s, 2H), 2.78 (s, 6H). **^13^C NMR**: (125 MHz, CD_3_OD:CD_3_CN 1:1, calibrated to CD_3_OD) δ (ppm) = 164.9, 138.3, 136.7, 135.6, 132.13, 132.07, 130.7, 126.4, 126.12, 126.08, 125.6, 116.9, 116.2, 113.3, 111.2, 42.0, 38.6. **HRMS**: (ESI) *m/z* calculated for C_18_H_20_N_4_O_3_S[M+H]^+^ 373.1329, found 373.1343.

### 2-amino-N-(3-(3-(N,N-dimethylsulfamoyl)phenyl)-1H-indol-5-yl)-2-methylpropanamide (18g)

The title compound was prepared according to general procedure B using the aryl-indole **44** (50 mg, 0.116 mmol), N-Boc-α-methyl alanine (24 mg, 0.116 mmol), PyBOP (601 mg, 0.116 mmol), DIPEA (0.15 mL, 0.58 mmol), and dry DMF (1.2 mL). RP-HPLC (gradient: 2-50 shortprep, tR = 8.50 min) of the crude deprotected product afforded the TFA salt of **18g** as a colorless solid (17 mg, 28%). **^1^H NMR**: (500 MHz, CD_3_CN) δ (ppm) = 9.75 (s, 1H), 8.75 (s, 1H), 8.20 (d, *J* = 1.9 Hz, 1H), 8.01 (dd, *J* = 1.7, 1.6 Hz, 1H), 7.95 (ddd, *J* = 7.3, 1.7, 1.6 Hz, 1H), 7.71 – 7.60 (m, 3H), 7.50 (d, *J* = 8.7 Hz, 1H), 7.34 (dd, *J* = 8.7, 1.9 Hz, 1H), 2.73 (s, 6H), 1.72 (s, 6H). **^13^C NMR**: (125 MHz, CD_3_CN) δ (ppm) = 170.6, 137.8, 136.5, 135.4, 131.9, 131.8, 130.6, 126.3, 125.9, 125.8, 125.6, 117.9, 116.2, 113.1, 112.2, 59.2, 38.6, 24.1. **HRMS**: (ESI) *m/z* calculated for C_20_H_24_N_4_O_3_S [M+H]^+^ 401.1642, found 401.1652.

### 1-amino-N-(3-(3-(N,N-dimethylsulfamoyl)phenyl)-1H-indol-5-yl)cyclopropane-1-carboxamide (18h)

The title com-pound was prepared according to general procedure B using the aryl-indole **44** (50 mg, 0.116 mmol), 1-((*tert*-butoxycarbonyl)amino)-cyclopropane-1-carboxylic acid (23 mg, 0.116 mmol), PyBOP (61 mg, 0.116 mmol), DIPEA (0.15 mL, 0.58 mmol), and dry DMF (1.16 mL). RP-HPLC (gradient: 2-50 shortprep, tR = 8.43 min) of the crude deprotected product afforded the TFA salt of **18h** as a colorless solid (16 mg, 27%). **^1^H NMR**: (500 MHz, CD_3_CN) δ (ppm) = 9.68 (s, 1H), 8.12 (d, *J* = 1.9 Hz, 1H), 8.01 (dd, *J* = 1.7, 1.7 Hz, 1H), 7.93 (ddd, *J* = 7.2, 1.8, 1.7 Hz, 1H), 7.89 (s, 1H), 7.70 – 7.62 (m, 3H), 7.49 (d, *J* = 8.7 Hz, 1H), 7.26 (dd, *J* = 8.8, 2.0 Hz, 1H), 2.74 (s, 6H), 1.74 – 1.68 (m, 2H), 1.60 – 1.54 (m, 2H). **^13^C NMR**: (125 MHz, CD_3_CN) δ (ppm) = 167.1, 136.9, 135.8, 134.6, 131.0, 130.3, 129.7, 125.4, 125.0, 124.9, 124.7, 118.2, 115.3, 112.2, 111.7, 37.6, 36.6, 12.4. **HRMS**: (ESI) *m/z* calculated for C_20_H_22_N_4_O_3_S [M+H]^+^ 399.1485, found 399.1495.

### 3-(5-(1,3-dioxoisoindolin-2-yl)-1H-indol-3-yl)-N,N-dimethylbenzenesulfonamide (45)

The title compound was prepared according to the general procedures I and F using **28** (64 mg, 0.12 mmol), TFA (1.2 mL), phthalic anhydride (18 mg, 0.12 mmol) and dry toluene (0.5 mL). Purification by column chromatography (EtOAc:hexanes 10:90) afforded **45** as a pale yellow solid (42 mg, 78%). **^1^H NMR**: (500 MHz, CDCl_3_) δ (ppm) = 8.58 (br s, 1H), 7.98 (d, *J* = 3.0 Hz, 1H), 7.96 (d, *J* = 3.0 Hz, 1H), 7.89 (br d, *J* = 8.5 Hz, 2H), 7.80 (d, *J* = 3.0 Hz, 1H), 7.79 (d, *J* = 3.0 Hz, 1H), 7.67 (br d, *J* = 7.7 Hz, 1H), 7.60-7.55 (m, 2H), 7.49 (br t, *J* = 2.2 Hz, 1H), 7.29 (br d, *J* = 8.8 Hz, 1H), 2.75 (s, 6H).

### 3-(5-(1,3-dioxoisoindolin-2-yl)-1-methyl-1H-indol-3-yl)-N,N-dimethylbenzenesulfonamide (46)

The title compound was prepared according to general procedure G using **45** (42 mg, 0.094 mmol), K_2_CO_3_ (65 mg, 0.47 mmol), MeI (0.018mL, 0.28 mmol) and dry DMF (0.5 mL). The crude product was directly used in the next step without further purification. **^1^H NMR**: (500 MHz, CDCl_3_) δ (ppm) = 7.97 (br s, 1H), 7.97 (d, *J* = 3.12 Hz, 1H), 7.96 (d, *J* = 3.0 Hz, 1H), 7.90 (br d, *J* = 1.9 Hz, 1H), 7.88 (ddd, *J* = 7.7, 1.5, 1.3 Hz, 1H), 7.80 (d, *J* = 3.0 Hz, 1H), 7.79 (d, *J* = 3.0 Hz, 1H), 7.65 (ddd, *J* = 7.9, 1.4, 1.3 Hz, 1H), 7.57 (dd, *J* = 7.9, 7.7 Hz, 1H), 7.50 (d, *J* = 8.7 Hz, 1H), 7.40 (s, 1H), 7.32 (dd, *J* = 8.7, 1.9 Hz, 1H), 3.91 (s, 3H), 2.75 (s, 6H).

### 3-(5-amino-1-methyl-1H-indol-3-yl)-N,N-dimethylbenzenesulfonamide (47)

The title compound was prepared according to general procedure H using **46** (37 mg, 0.080 mmol), hydrazine hydrate (3.4 mg, 0.10 mmol) and MeOH (1.3 mL). The crude product was directly used in the next step without further purification. **^1^H NMR**: (500 MHz, CDCl_3_) δ (ppm) = 7.99 (br t, *J* = 1.7 Hz, 1H), 7.85 (dt, *J* = 7.5, 1.5 Hz, 1H), 7.60 (dt, *J* = 8.0, 1.7 Hz, 1H), 7.55 (dd, *J* = 17.2, 7.8 Hz, 1H), 7.25 (s, 1H), 7.21 (d, *J* = 2.2 Hz, 1H), 7.19 (d, *J* = 8.5 Hz, 1H), 6.77 (dd, *J* = 8.5, 2.2 Hz, 1H), 3.80 (s, 3H), 2.75 (s, 3H).

### (S)-2-amino-N-(3-(3-(N,N-dimethylsulfamoyl)phenyl)-1-methyl-1H-indol-5-yl)propanamide (20)

The title com-pound was prepared according to general procedure B using the aryl-indole **47** (28 mg, 0.080 mmol), (*tert*-butoxycarbonyl)*-L*-alanine (18 mg, 0.095 mmol), PyBOP (50 mg, 0.38 mmol), DIPEA (0.070 mL, 0.095 mmol), and dry DMF (0.8 mL) followed by Boc-deprotection using TFA (0.7 mL). Purification by RP-HPLC (using a SiliCycle SiliaChrom dtC18 semipreparative column (5 μm, 100À, 10 x 250 mm) with a flow rate of 5 mL/min eluting with solvent (A: 0.1 % TFA in water B: 0.1 % TFA in MeCN) on a gradient of (2 → 100) %) solvent B over 15 min, t_R_ = 6.5 min) of the crude deprotected product afforded the TFA salt of **20** as a colorless solid (10 mg, 15%). **^1^H NMR**: (400 MHz, CD_3_CN) δ (ppm) = 9.23 (s, 1 H), 8.23 (d, *J* = 1.6 Hz, 1H), 7.96 (br s, 1H), 7.87 (dt, *J* = 7.4, 1.6 Hz, 1H), 7.63 (t, *J* = 7.4 Hz, 1H), 7.60 (t, *J* = 1.6 Hz, 1H), 7.55 (s, 1H), 7.35 (dq, *J* = 8.9, 0.8 Hz, 1H), 7.29 (dd, *J* = 8.9, 1.9 Hz, 1H), 4.24 (q, *J* = 7.0 Hz, 1H), 3.79 (s, 3H), 2.71 (s, 6H), 1.58 (d, *J* = 7.0 Hz, 3H). **^13^C NMR**: (150 MHz, CD_3_CN) δ (ppm) = 168.6, 137.7, 136.6, 136.0, 132.2, 131.6, 130.6, 130.2, 126.2, 126.1, 125.4, 116.8, 114.9, 111.4, 111.3, 50.96, 38.60, 33.47, 17.62. **HRMS**: (ESI) *m/z* calculated for C_21_H_2_4N4O_3_S [M+H]^+^ 401.1647, found 401.1602.

### ((benzyloxy)carbonyl)-L-serine (48)

1.17 g NaHCO_3_ (14 mmol, 1 equiv.) and 3.45 g Na_2_CO_3_ (28 mmol, 2 equiv.) were dissolved in 3 mL acetone and 32 mL H_2_O. *L*-serine (1.46 g, 14 mmol, 1 equiv.) was then added, followed by CbzCl (2.47 mL, 17 mmol, 1.25 equiv.). The mixture was then stirred vigorously at room temperature. Each time the mixture became cloudy, 1M KOH was added until it became a clear solution again. After 5h, the reaction mixture was washed three times with Et2O and the aqueous layer was acidified with concentrated HCl. Filtration afforded the title compound as a white powder (2.38 g, 72%). **^1^H NMR:** (400 MHz, DMSO-d_6_) δ (ppm) = 12.63 (s, 1H), 7.42 – 7.27 (m, 5H), 5.05 (s, 2H), 4.11 – 4.02 (m, 1H), 3.72 – 3.60 (m, 2H).

### N-((benzyloxy)carbonyl)-O-(tert-butyldimethylsilyl)-L-serine (49)

To a suspension of **48** (2.38 g, 10 mmol, 1 equiv.) and imidazole (2.03 g, 30 mmol, 3 equiv.) in CH_2_Cl_2_ (56 mL) were added TBSCl (1.65 g, 11 mmol, 1.1 equiv.) and DMAP (61 mg, 0.50 mmol, 0.05 equiv.) and the resulting solution was then stirred at room temperature overnight. The reaction mixture was then diluted with 1M HCl and H_2_O and the layers were separated. The aqueous were extracted twice more with CH_2_Cl_2_ and the combined organic were washed with 1M HCl, H_2_O and brine, dried over Na_2_SO_4_, filtered and evaporated to afford a crude oil. This material was redissolved in MeOH/PhMe and the solvents were evaporated again. This process of MeOH/PhMe addition/evaporation was repeated twice more to afford the title compound as a thick white oil (2.81 g, 80%). **^1^H NMR:** (400 MHz, CDCl_3_) δ (ppm) = 7.45 – 7.29 (m, 5H), 5.58 (d, *J* = 8.2 Hz, 1H), 5.19 – 5.05 (m, 2H), 4.44 (t, *J* = 5.8 Hz, 1H), 4.17 – 4.05 (m, 1H), 3.85 (td, *J* = 9.9, 3.5 Hz, 1H), 0.86 (s, 9H), 0.03 (s, 6H).

### benzyl (S)-4-(((tert-butyldimethylsilyl)oxy)methyl)-5-oxooxazolidine-3-carboxylate (50)

A flask was charged with **49** (2.81 g, 8 mmol, 1 equiv.), pTsOH.H_2_O (227 mg, 1.2 mmol, 0.15 equiv.), (CH_2_O)_n_ (1.40 g) and dry PhMe (115 mL). A Dean-Stark apparatus was attached and the mixture was heated to reflux for 1h. The reaction mixture was then washed with saturated NaHCO_3_, H_2_O and brine, dried over Na_2_SO_4_, filtered and evaporated to afford 4.34 g of a crude yellow oil. Purification by flash column chromatography (EtOAc :hexanes 30:70) afforded the title compound as a colourless oil (1.81 g, 62%). **^1^H NMR:** (400 MHz, CDCl_3_) δ (ppm) = 7.43 – 7.30 (m, 5H), 5.53 (d, *J* = 27.9 Hz, 1H), 5.22 (s, 2H), 4.38 – 4.16 (m, 3H), 4.05 – 3.98 (m, 1H), 0.84 (s, 9H), 0.10 (s, 3H), 0.01 (s, 3H).

### benzyl (R)-4-(fluoromethyl)-5-oxooxazolidine-3-carboxylate (51)

To a solution of **50** (1.81 g, 5 mmol, 1 equiv.) in dry CH_2_Cl_2_ (25mL) at −78ºC were added Et_3_N.HF (3.23 mL, 20 mmol, 4 equiv.) and Xtalfluor-E (2.27 g, 10 mmol, 2 equiv.). The mixture was stirred for 1h at −78 °C and then allowed to warm to room temperature overnight. The reaction mixture was then diluted with saturated NaHCO_3_ and stirred vigorously for 20min. The layers were then separated and the aqueous was extracted twice more with CH_2_Cl_2_. The combined organic were washed with saturated NaHCO_3_, H_2_O and brine, dried over Na_2_SO_4_, filtered and evaporated to afford 847mg of a brown oil. Purification by flash column chromatography (EtOAc:hexanes 30:70) afforded the title compound as a pale yellow oil (442 mg, 35%). **^1^H NMR:** (400 MHz, CDCl_3_) δ (ppm) =7.44 – 7.32 (m, 5H), 5.55 (s, 1H), 5.36 – 5.15 (m, 3H), 4.83 (dd, *J* = 45.0, 9.3 Hz, 2H), 4.39 (d, *J* = 33.9 Hz, 1H). ^**19**^F NMR (376 MHz, CDCl_3_) δ (ppm) = (−231.56 – (−232.71)) (m, 1F).

### (R)-2-((tert-butoxycarbonyl)amino)-3-fluoropropanoic acid (52)

To a solution of **51** (241 mg, 0.95 mmol, 1 equiv.) in dry CH_2_Cl_2_ (8 mL) at 0 °C was added 1M BCl_3_ in CH_2_Cl_2_ (3.81 mL, 3.81 mmol, 4 equiv.) dropwise. The ice bath was removed and the mixture was allowed to warm to room temperature over 25 min. The reaction mixture was then poured into 25 mL ice-water and the layers were separated. The aqueous layer was washed once with CH_2_Cl_2_ and then it was neutralized with 5M KOH. Evaporation of the water afforded 1.06 g of a white solid, which was redissolved in 3 mL H_2_O and cooled to 0 °C. To that solution, NaOH (57 mg, 1.43 mmol, 1.5 equiv.) was added, followed by THF (3 mL). Finally, Boc_2_O (270 mg, 1.24 mmol, 1.3 equiv.) was also added and the mixture was allowed to warm to room temperature overnight. The reaction mixture was then washed twice with hexanes and the aqueous layer was acidified carefully with 1M HCl until pH = 1. Then, the aqueous were extracted four times with EtOAc and the combined organic extracts were dried over Na_2_SO_4_, filtered and evaporated to afford 132 mg of an amber oil. This material was dissolved in MeOH/EtOAc and filtered through a silica plug to afford the title compound as a yellow oil (98 mg, 50%). **^1^H NMR:** (400 MHz, CD_3_OD) δ (ppm) = 4.72 – 4.65 (m, 1H), 4.57 (dd, *J* = 9.4, 3.2 Hz, 1H), 4.39 (d, 29.8 Hz, 2H), 1.46 (s, 9H). ^**19**^F NMR (376 MHz, CD_3_OD) δ (ppm) −230.19 (s, 1F).

### tert-butyl (R)-(1-((3-(3-(N,N-dimethylsulfamoyl)phenyl)-1-methyl-1H-indol-5-yl)amino)-3-fluoro-1-oxopropan-2-yl)carbamate (53)

A vial was charged with **47** (11 mg, 0.03 mmol, 1 equiv.) and **52** (14 mg, 0.07 mmol, 2 equiv.). Then, dry CH_2_Cl_2_ (250 μL) was added, followed by DIPEA (35 μL, 0.13 mmol, 4 equiv.) and HATU (27 mg, 0.07 mmol, 2.1 equiv.). The reaction mixture was then stirred at room temperature overnight. It was then washed with 1M HCl and brine, dried over Na_2_SO_4_, filtered and evaporated to afford 32 mg of a brown oil. Purification by flash column chromatography (EtOAc :hexanes 3:2) afforded the title compound as a yellow oil (12 mg, 69%). **^1^H NMR:** (400 MHz, CD_3_CN) δ (ppm) = 8.65 (s, 1H), 8.23 (d, *J* = 1.9 Hz, 1H), 8.02 – 7.97 (m, 1H), 7.93 (tt, *J* = 7.5, 1.6 Hz, 1H), 7.71 – 7.61 (m, 1H), 7.65 – 7.56 (m, 2H), 7.45 – 7.39 (m, 1H), 7.39 – 7.30 (m, 1H), 4.85 – 4.59 (m, 2H), 4.45 (d, *J* = 12.6 Hz, 1H), 3.84 (d, *J* = 3.2 Hz, 3H), 2.72 (s, 6H), 1.45 (s, 9H). ^**19**^F NMR: (376 MHz, CD_3_CN) δ (ppm) = −229.00 (s, 1F).

### (R)-2-amino-N-(3-(3-(N,N-dimethylsulfamoyl)phenyl)-1-methyl-1H-indol-5-yl)-3-fluoropropanamide (21)

To a so-lution of **53** (12 mg, 0.02 mmol, 1 equiv.) in dry CH_2_Cl_2_ (200 μL), TFA (200 μL, 2.6 mmol, 131 equiv.) was added. After 1h at room temperature, the solvents were evaporated. Purification by RP-HPLC (using a SiliCycle SiliaChrom dtC18 semipreparative column (5 μm, 100À, 10 x 250 mm) with a flow rate of 5 mL/min eluting with solvent (A: 0.1 % TFA in water B: 0.1 % TFA in MeCN) on a gradient of (2 → 100) %) solvent B over 15 min, t_R_ = 6.60 min) afforded the TFA salt of **21** as a colourless solid (3 mg, 24%). **^1^H NMR:** (400 MHz, CD_3_CN) δ (ppm) = 9.44 (s, 1H), 8.24 (s, 1H), 7.98 (s, 1H), 7.91 – 7.89 (m, 1H), 7.66 – 7.58 (m, 3H), 7.40 (d, *J* = 8.7 Hz, 1H), 7.32 (d, *J* = 8.8 Hz, 1H), 4.96 (d, *J* = 46.5 Hz, 2H), 4.46 (s, 1H), 3.82 (s, 3H), 2.72 (s, 6H). ^**19**^F NMR: (376 MHz, CD_3_CN) δ (ppm) = −231.3 (s, 1F). **HRMS:** (ESI) *m/z* calculated for C_20_H_23_FN_4_O_3_S [M+H]^+^ 419.1548, found 419.1565.

### 3-bromo-N,N-dimethylbenzamide (54)

To a solution of 2M Me_2_NH in THF (11.4 mL, 23 mmol, 2 equiv.) in 25 mL dry CH_2_Cl_2_ at 0 °C, *m*-bromobenzoyl chloride (1 mL, 7.6 mmol, 1 equiv.) was added dropwise. After the addition was complete, the reaction mixture was allowed to warm to room temperature overnight. After reaction was completed as observed by TLC, it was diluted with saturated NaHCO_3_. The layers were separated and the aqueous was extracted once more with CH_2_Cl_2_. The combined organic layers were dried over Na_2_SO_4_, filtered and evaporated to afford the title compound as a red oil (1.73 g, quantitative). **^1^H NMR:** (500 MHz, CDCl_3_) δ 7.53 (t, *J* = 1.7 Hz, 1H), 7.51 (ddd, *J* = 7.9, 2.0, 1.2 Hz, 1H), 7.31 (dt, *J* = 7.7, 1.3 Hz, 1H), 7.25 – 7.22 (m, 1H), 3.01 (d, *J* = 64.7 Hz, 6H).

### N,N-dimethyl-3-(4,4,5,5-tetramethyl-1,3,2-dioxaborolan-2-yl)benzamide (55)

A vial was purged with N_2_ and charged with **54** (765 mg, 3.4 mmol, 1 equiv.), B_2_pin_2_ (937 mg, 3.7 mmol, 1.1 equiv.), KOAc (1.65 g, 17 mmol, 5 equiv.), dioxane (15 mL) and H_2_O (2 mL). The seal was removed temporarily and Pd(dppf)Cl_2_.CH_2_Cl_2_ (137 mg, 0.17 mmol, 5 mol%) was added. The vial was then sealed again and microwaved for 2h at 120 °C. The reaction mixture was then diluted with saturated NaHCO_3_ and brine and extracted with EtOAc (3 times). The combined organic layers were then washed with brine, dried over Na_2_SO_4_, filtered and evaporated to afford a brown oil (977mg). Purification by column chromatography (EtOAc:hexanes 3:1) afforded a yellow oil, which was a 1:1.7 mixture of **55**:B_2_pin_2_ (487 mg, 21%). This material was used for the next step without further purification. **^1^H NMR:** (400 MHz, CDCl_3_) δ 7.88 – 7.78 (m, 2H), 7.53 – 7.47 (m, 1H), 7.39 (td, *J* = 7.5, 0.8 Hz, 1H), 3.05 (d, *J* = 52.9 Hz, 6H), 1.34 (s, 12H).

### tert-butyl 5-((tert-butoxycarbonyl)amino)-3-(3-(dimethylcarbamoyl)phenyl)-1H-indole-1-carboxylate (56)

The title compound was prepared according to general procedure A using **7** (105 mg, 0.26 mmol, 1equiv.), 3-pinacolboronate arylbenzamide **55** mixture (77 mg, 0.28 mmol, 1.1 equiv.) K_2_CO_3_ (106 mg, 0.77 mmol, 3 equiv.), Pd(dppf)Cl_2_.CH_2_Cl_2_ (21 mg, 0.03 mmol, 0.1 equiv.), degassed dry THF (3.47 mL) and degassed H_2_O (1.13 mL). Purification by column chromatography (EtOAc:hexanes 20:80 → 66:33) afforded the title compound as a brown oil (81 mg, 66%). **^1^H NMR:** (400 MHz, CDCl_3_) δ (ppm) = 8.12 (d, *J* = 9.0 Hz, 1H), 7.86 (s, 1H), 7.76 – 7.58 (m, 3H), 7.58 – 7.43 (m, 1H), 7.44 – 7.35 (m, 1H), 7.29 (d, *J* = 9.2 Hz, 1H), 6.57 (s, 1H), 3.08 (d, *J* = 26.3 Hz, 6H), 1.68 (s, 9H), 1.52 (s, 9H).

### 3-(5-(1,3-dioxoisoindolin-2-yl)-1-methyl-1H-indol-3-yl)-N, N-dimethylbenzamide (57)

The title compound was prepared according to the general procedures I, F and G using **56** (60 mg, 0.12 mmol), TFA (1.2 mL), phthalic anhydride (18 mg, 0.12 mmol), dry toluene (0.25 mL), K_2_CO_3_ (86 mg, 0.62 mmol), MeI (0.024 mL, 0.40 mmol) and dry DMF(0.63 mL). Purification by column chromatography (MeOH:CH_2_Cl_2_ 10:90) afforded **56** as a pale yellow solid (42 mg, 85%). **^1^H NMR**: (500 MHz, CDCl_3_) δ (ppm) = 8.01 (s, 2H), 7.96 (d, *J* = 3.1 Hz, 1H), 7.95 (d, *J* = 3.1 Hz, 1H), 7.88 (d, *J* = 1.8 Hz, 1H), 7.79 (d, *J* = 3.1 Hz, 1H), 7.78 (d, *J* = 3.1 Hz, 1H), 7.67 (dt, *J* = 7.7, 1.4 Hz, 1H), 7.62 (t, *J* = 1.4 Hz, 1H), 7.48 (d, *J* = 8.7 Hz, 1H), 7.43 (t, *J* = 7.7 Hz, 1H), 7.32 (s, 1H), 7.29 (dt, *J* = 7.4, 1.3 Hz, 1H), 3.88 (s, 3H), 3.12 (s, 3H), 3.02 (s, 3H).

### 3-(5-amino-1-methyl-1H-indol-3-yl)-N,N-dimethylbenzamide (58)

The title compound was prepared according to general procedure H using **57** (42 mg, 0.10 mmol), hydrazine hydrate (7 mg, 0.14 mmol) and MeOH (1.0 mL). The crude product was directly used in the next step without further purification. **^1^H NMR**: (500 MHz, CDCl_3_) δ (ppm) = 7.64 (d, *J* = 1.5 Hz, 1H), 7.63 (ddd, *J* = 4.4, 1.8, 1.4 Hz, 1H), 7.42 (t, *J* = 7.6 Hz, 1H), 7.26 (s, 1H), 7.25 (dt, *J* = 7.1, 1.4 Hz, 1H), 7.23 (d, *J* = 2.2 Hz, 1H), 7.16 (d, *J* = 7.6 Hz, 1H), 6.74 (dd, *J* = 8.6, 2.2 Hz, 1H), 3.76 (s, 3H), 3.13 (br s, 3H), 3.02 (br s, 3H).

### (S)-3-(5-(2-aminopropanamido)-1-methyl-1H-indol-3-yl) - N,N-dimethylbenzamide (22)

The title compound was prepared according to general procedure B using the aryl-indole **58** (14 mg, 0.048 mmol), (*tert*-butoxycarbonyl)*-L*-alanine (11 mg, 0.057 mmol), PyBOP (30 mg, 0.24 mmol), DIPEA (0.041 mL, 0.057 mmol), and dry DMF (0.5 mL) followed by Boc-deprotection using TFA (0.5 mL). Purification by RP-HPLC (using a SiliCycle SiliaChrom dtC18 semipreparative column (5 μm, 100À, 10 x 250 mm) with a flow rate of 5 mL/min eluting with solvent (A: 0.1 % TFA in water B: 0.1 % TFA in MeCN) on a gradient of (2 → 100) %) solvent B over 15 min, t_R_ = 5.95 min) of the crude deprotected product afforded the TFA salt of **22** as a colorless solid (7 mg, 40%). **^1^H NMR**: (400 MHz, CD_3_CN) δ (ppm) = 9.10 (s, 1H), 8.22 (s, 1H), 7.67 (d, *J* = 7.3, 2H), 7.49 (s, 1H), 7.46 (t, *J* = 7.9 Hz, 1H), 7.38 (d, *J* = 8.8 Hz, 1H), 7.29 (br d, *J* = 8.8 Hz, 1H), 7.25 (br d, *J* = 7.6 Hz, 1H), 4.19 (q, *J* = 6.4 Hz, 1H), 3.80 (s, 3H), 3.05 (br s, 3H), 2.99 (br s, 3H), 1.58 (d, *J* = 6.4 Hz, 3H). **^13^C NMR**: (150 MHz, CD3CN) δ (ppm) = 172.1, 168.6, 138.4, 136.7, 135.9, 131.9, 129.8, 129.6, 128.5, 126.4, 126.0, 125.0, 116.6, 116.0, 111.6, 111.3, 51.02, 39.89, 35.39, 33.50, 30.20, 17.57. **HRMS**: (ESI) *m/z* calculated for C_21_H_24_N_4_O_2_ [M+H]^+^ 365.4565, found 365.1915.

## ASSOCIATED CONTENT

**Supporting Information**. The supporting information is available free of charge at http://pubs.acs.org.

## ACKNOWLEDGMENT

This work was supported by a Natural Sciences and Engineering Research Council (NSERC) Discovery Grant (RB 2019-06368), and NSERC CREATE Grant (432008-2013). The SGC is a registered charity (Number 1097737) that receives funds from AbbVie, Bayer Pharma AG, Boehringer Ingelheim, Canada Foundation for Innovation, Eshelman Institute for Innovation, Genome Canada through Ontario Genomics Institute [OGI-055], Innovative Medicines Initia-tive (EU/ EFPIA) [ULTRA-DD Grant No. 115766], Janssen, Merck KGaA, Darmstadt, Germany, MSD, Novartis Pharma AG, Ontario Ministry of Research, Innovation and Science (MRIS), Pfizer, Sâo Paulo Research Foundation-FAPESP, Takeda, and Wellcome [106169/ZZ14/Z].

## ABBREVIATIONS

PRMT4: Protein Arginine Methyl Transferase 4
PRMT6: Pro-tein Arginine Methyl Transferase 6
IC_50_: half maximal inhibi-tory concentration
Boc: tert-butyloxycarbonyl
NBS: N-bro-mosuccinimide
pyBOP: Benzotriazole-1-yl-oxy-tris-pyrrolidino-phosphonium hexafluorophosphate
B2pin2: Bis(pinacolato)diboron
BAF155: SMARCC1 gene protein

